# Functionally selective dopamine D1 receptor endocytosis and signaling by catechol and non-catechol agonists

**DOI:** 10.1101/2024.04.15.589637

**Authors:** Ashley N. Nilson, Daniel E. Felsing, Pingyuan Wang, Manish K. Jain, Jia Zhou, John A. Allen

**Author notes:** **Corresponding author**: John A. Allen, Ph.D., Assistant Professor, Department of Pharmacology and Toxicology, Center for Addiction Research, School of Medicine, University of Texas Medical Branch, 301 University Boulevard, MC 0615, Bldg 17 3.324 Galveston, TX, 77555-0615, Office –. Primary laboratory of origin: John A. Allen Ph.D., Assistant Professor, Department of Pharmacology and Toxicology, University of Texas Medical Branch, Galveston, Texas 77555-0615, United States.

## Abstract

The dopamine D1 receptor (D1R) has fundamental roles in voluntary movement and memory and is a validated drug target for neurodegenerative and neuropsychiatric disorders. However, previously developed D1R selective agonists possess a catechol moiety which displays poor pharmacokinetic properties. The first selective non-catechol D1R agonists were recently discovered and unexpectedly many of these ligands showed G protein biased signaling. Here, we investigate both catechol and non-catechol D1R agonists to validate potential biased signaling and examine if this impacts agonist-induced D1R endocytosis. We determined that most, but not all, non-catechol agonists display G protein biased signaling at the D1R and have reduced or absent β-arrestin recruitment. A notable exception was compound (Cmpd) 19, a non-catechol agonist with full efficacy at both D1R-G protein or D1R-β-arrestin pathways. In addition, the catechol ligand A-77636 was a highly potent, super agonist for D1R-β-arrestin activity. When examined for agonist-induced D1R endocytosis, balanced agonists SKF-81297 and Cmpd 19 induced robust D1R endocytosis while the G protein biased agonists did not. The β-arrestin super agonist, A-77636, showed significantly increased D1R endocytosis. Moreover, β-arrestin recruitment efficacy of tested agonists strongly correlated with total D1R endocytosis. Taken together, these results indicate the degree of D1R signaling functional selectivity profoundly impacts D1R endocytosis regardless of pharmacophore. The range of functional selectivity of these D1R agonists will provide valuable tools to further investigate D1R signaling, trafficking and therapeutic potential.

**Significance Statement:** The D1R is a validated therapeutic target and the recently discovered non-catechol D1R agonists have translational potential. We have systematically characterized several structurally distinct D1R agonists including non-catechols with balanced or G protein biased activity. When examined for agonist-induced D1R endocytosis, balanced agonists induced robust D1R endocytosis while G protein biased agonists did not. These results indicate the degree of D1R signaling functional selectivity profoundly impacts receptor endocytosis. This work also independently validates agonist tools to further investigate D1R activation in basic and translational research.

## Introduction

The dopamine D1 receptor (D1R) is a G protein-coupled receptor (GPCR) that regulates voluntary movement, working memory, attention and reward (Arnsten et al., 2017; Beaulieu and Gainetdinov, 2011; Dichter et al., 2012; Girgis et al., 2016; Goldman-Rakic et al., 2004; Nutt et al., 2015). The D1R is a validated drug target with the potential to treat motor deficits in Parkinson’s disease, impaired working memory, and reward pathway dysfunction in neuropsychiatric disorders (Arnsten et al., 2017; Beaulieu and Gainetdinov, 2011; Dichter et al., 2012; Nilson et al., 2022). The D1R couples to Gs/Golf G proteins that activate adenylyl cyclase to stimulate cyclic adenosine monophosphate (cAMP) production (Beaulieu and Gainetdinov, 2011). The related dopamine D5 receptor (D5R) is a highly homologous receptor that functions similarly to the D1R. In addition, agonist activation of the D1R recruits the multi-functional adaptor protein β-arrestin to the receptor, which can induce D1R desensitization and receptor endocytosis (Beaulieu and Gainetdinov, 2011; Kim et al., 2004; Nilson et al., 2022; Vickery and von Zastrow, 1999), which may support continued D1R-G protein signaling at endosomes (Irannejad et al., 2013; Kotowski et al., 2011).

Selectively targeting the D1R holds great potential to alleviate the symptoms of several neurological and neuropsychiatric disorders; however, previous efforts have largely failed in developing drug-like D1R agonists for clinical use due to pharmacokinetic limitations (Felsing et al., 2019; Waddington, 1988). Until recently, all D1R selective agonists shared a common chemical structure containing a dihydroxyphenyl catechol moiety that is also present in catecholamine neurotransmitters (Felsing et al., 2019). Catechol D1R agonists have poor oral bioavailability, are rapidly metabolized and in some cases cause drug-induced tolerance (Asin and Wirtshafter, 1993; Blanchet et al., 1998; Gray et al., 2018; Wade and Nomikos, 2005). Recently, Pfizer made a breakthrough discovery of the first non-catechol D1R selective agonists that have drug-like pharmacokinetics allowing sustained brain exposure and the compounds have shown promising results in clinical trials for Parkinson’s disease (Coe JW et al., 2014; Davoren, 2015; Gray et al., 2018; Riesenberg et al., 2020; Sohur et al., 2018). The discovery of non-catechol D1R agonists provides tools with unprecedented pharmacokinetics. Notably, the reported non-catechol agonists showed functionally selective D1R-G protein signaling with reduced recruitment of β-arrestin (Gray et al., 2018). Recently, analogs of the non-catechol agonists have been synthesized that show varying degrees of D1R functional selectivity (Davoren et al., 2018; Martini et al., 2019a; Martini et al., 2019b).

It is now well-established that functionally selective agonists, also known as biased agonists, exist for many GPCRs (Urban et al., 2007). Biased agonists may fine tune therapeutic activity improving efficacy or reducing side effects (Allen et al., 2011; Gray et al., 2018; Schmid et al., 2017; Smith et al., 2018). However, the effects of D1R signaling *in vitro* and *in vivo* by biased agonists remains largely undefined. For example, it is currently unclear if the G protein biased D1R agonists, which have reduced β-arrestin recruitment, also influence D1R endocytosis differentially than balanced agonists. Furthermore, the therapeutic utility of targeting one pathway over the other (G protein versus β-arrestin) has not been fully defined at the D1R. Functionally selective agonists that preferentially activate D1R-G protein signaling or D1R-β-arrestin signaling will be valuable tools to further define D1R molecular functions and evaluate the therapeutic utility of biased D1R activation (Arnsten et al., 2017; Conroy et al., 2015; Felsing et al., 2019; Yang et al., 2018)

Here, we pursue pharmacological studies testing both catechol and non-catechol agonists with a primary goal to determine if agonist functional selectivity impacts D1R endocytosis. We determined that most, but not all, non-catechol agonists are G protein biased at the D1R and have reduced or absent β-arrestin recruitment. A notable exception was compound (Cmpd) 19, a non-catechol agonist with full efficacy at both D1R-G protein and D1R-β-arrestin pathways. The G protein biased functional selectivity was also similar when ligands were tested for agonist activity at the closely related D5R. In addition, the catechol ligand A-77636 was a highly potent, super agonist for D1R-β-arrestin activity. When examined for agonist-induced D1R endocytosis, balanced agonists SKF-81297 and Cmpd 19 induced robust D1R endocytosis while the G protein biased agonists did not. The β-arrestin super agonist, A-77636, also showed the strongest ability to induce D1R endocytosis. Moreover, β-arrestin recruitment efficacy of agonists strongly correlated with total D1R endocytosis. Together these results indicate the degree of functional selectivity, regardless of pharmacophore, extends to profoundly impact D1R endocytosis. These balanced and biased agonist tools will be useful to further investigate D1R signaling, membrane trafficking and therapeutic potential.

## Materials and Methods

### Materials

The commercially available compounds were purchased from the following suppliers: dopamine (Sigma, St. Louis, MO), SKF-81297 (Sigma, St. Louis, MO), SKF-38393 (Tocris, Bristol, UK), SKF-77434 (Tocris, Bristol, UK), A-77636 (Tocris, Bristol, UK), ascorbic acid (Sigma, St. Louis, MO). D1R non-catechol agonists, which are not commercially available, were synthesized at the University of Texas Medical Branch by Dr. Jia Zhou’s laboratory. PF-1119, PF-2334, PF-6142 were resynthesized according to protocols described in Gray et al. (2018). 6-(4-(Furo[3,2-*c*]pyridin-4-yloxy)-2-methylphenyl)-1,5-dimethylpyrimidine-2,4(1*H*,3*H*)-dione (Cmpd 19) was resynthesized and was originally disclosed and structure described by Pfizer, Inc. in International Patent Publication No. WO2014/072881(Coe JW et al., 2014) and the subsequently approved US Patent 96172751(Coe JW et al., 2017). Some compounds under study in this report were previously described by our group in a research letter as being either biased or unbiased D1R agonists, including a described synthesis and the balanced activity of Cmpd 19 (Wang et al., 2019). However, the letter describing Cmpd 19 was subsequently retracted (Wang et al., 2022). Therefore, we wish to confirm that all results described in this current report are original and unpublished findings, including the described activity of Cmpd 19 herein. 6-(4-(Furo[3,2-*c*]pyridin-4-yloxy)-2-methylphenyl)-5-methylpyrazin-2-amine (Cmpd 7), 4-(4-(Imidazo[1,2-*a*]pyridin-5-yl)-3-(trifluoromethyl)phenoxy)-furo[3,2-*c*]pyridine (Cmpd 41), and 7-(4-(Imidazo[1,2-*a*]pyridin-5-yl)-3-(trifluoromethyl)phenoxy)-thieno[2,3-*c*]pyridine (Cmpd 46) were resynthesized according to synthetic protocols described by Martini et al. (2019a). All synthesized compounds were verified for purity (>98%) by HPLC analysis and chemical structures verified using NMR and MS analytical techniques.

### DNA constructs

All DNA plasmid constructs used were in a pcDNA3.1 plasmid backbone. N-terminal 3X hemagglutinin (HA) tagged human D1R (HA-D1R) plasmid was purchased from cDNA Resource Center (Catalog #DRD01TN00, cDNA Resource Center Bloomsburg University, Bloomsburg, PA). DRD1-TANGO or DRD5-TANGO plasmid DNA were created by Bryan Roth and obtained from Addgene (Plasmids #66268 or #66272, Addgene, Watertown, MA). The β-arrestin2 green fluorescent protein (GFP) plasmid DNA was originally created by Robert Lefkowitz and obtained from Addgene (plasmid #35411, Addgene, Watertown, MA;). The 22F-Glosensor plasmid was purchased from Promega, Madison, WI.

The D5R-STOP construct used for D5R cAMP assays was made by site-directed mutagenesis to remove the engineered Tango C-terminal tail from the D5R-TANGO construct. Point mutation of D5R-STOP from D5-TANGO was made by polymerase chain reaction (PCR) using the QuikChange II Site–Directed Mutagenesis Kit (Agilent Technologies, Santa Clara, CA) according to the manufacturer’s protocol. Mutagenesis and sequencing primers were obtained from ThermoFisher (Pittsburgh, PA). The primers used to make the D5R-STOP point mutant were 5’– cgcccaacggatttcactaagataccggtggacgcac–3’ (sense) and 5’– gtgcgtccaccggtatcttagtgaaatccgttgggcg –3’ (antisense). Parental DNA in the reaction mixture was digested using restriction endonuclease DpnI (from diplococcus pneumoniae) at 37°C for one hour. Digestion mixture (2 µL) was transformed into XL1– Blue supercompetent cells by heat shock at 42°C for 45 seconds. The reaction was incubated in Super Optimal broth with Catabolite repression medium (Sigma–Aldrich) and then plated onto Luria–Bertani (LB) agar plates containing 100 µg/mL ampicillin. A single colony was selected for sequencing, and the point mutation verified prior to use (Molecular Genomics Core, University of Texas Medical Branch, Galveston, TX).

### Cells and Cell Culture

Human embryonic kidney 293 (HEK293) cells stably expressing the human D1R (D1-HEK) (up to pass 35) were maintained in Dulbecco’s modified Eagle’s medium (DMEM, Gibco/Fisher Scientific, Hampton, NH) supplemented with 10% fetal bovine serum (Omega Scientific, Tarzana, CA), 1% penicillin/streptomycin (Gibco/Fisher Scientific, Hampton, NH), 1% minimum essential medium non-essential amino acids (Gibco/Fisher Scientific, Hampton, NH), 25 mM HEPES (Gibco/Fisher Scientific, Hampton, NH), and 500 ug/ml Geneticin (Gibco/Fisher Scientific, Hampton, NH). HTLA cells (HEK293 cells stably expressing tTA-dependent luciferase reporter and a β-arrestin2-TEV fusion gene) (up to pass 32) were described previously (Kroeze et al., 2015) and generously provided by Dr. Bryan Roth (University of North Carolina at Chapel Hill) and maintained in DMEM supplemented with 10% fetal bovine serum, 1% penicillin/streptomycin, 2 µg/mL Puromycin (Gibco/Fisher Scientific, Hampton, NH), and 100 µg/mL Hygromycin (Thermofisher, Waltham, MA). Unmodified HEK293 cells (up to pass 25) were maintained in DMEM supplemented with 10% fetal bovine serum and 1% penicillin/streptomycin. All cell culturing was conducted in a humidified 37° C cell culture incubator with 5% CO_2_.

### D1R/D5R cAMP assay

HEK293 cells stably expressing the human D1R (D1-HEK) were split into poly-L-lysine (R&D systems, Minneapolis, MN) coated 6-well plates at 450,000 cells per well. After 24 hours incubation, the HEK293 cells were transfected using 1.0 µg 22F-Glosensor plasmid (Promega, Madison, WI) and 10 ul Lipofectamine2000 (Invitrogen, Carlsbad, CA) per well of a 6-well plate as per manufacturer’s protocol. The D1-HEK cells were transfected overnight and the following morning split into poly-L-lysine (R&D systems, Minneapolis, MN) coated 96-well white clear bottom cell culture plates (Greiner Bio-One, Monroe, MS) at 50,000 cells per well. Approximately 48 hours after transfection, the media was discarded and Glosensor substrate reagent (Promega, Madison, WI) was diluted to 1% in 20 mM HEPES in 1X Hank’s balanced salt solution (HBSS, Gibco/Fisher Scientific, Hampton, NH) and 90 µL of reagent was added to cells and incubated for 2 hours at room temperature in the dark. Agonist concentration treatments consisted of 0.1% dimethyl sulfoxide (DMSO) for basal cAMP counts or eleven concentrations (10 pM to 10 µM) of test compound. 10 µL of compound at 10x final concentration were added to wells containing 90 µL of Glosensor reagent and the cells were incubated 10 minutes at room temperature. Agonist response (light counts per second) was measured using a Microbeta2 plate reader (Perkin Elmer, Waltham, MA), capturing the total light counts in each well in one second. The highest concentration of DMSO was 0.1% DMSO at 10 µM except where noted below, and this was used as the vehicle control for basal cAMP measurements. We observed that Cmpd 7 and Cmpd 46 were not soluble at high concentrations unlike all other tested agonists. Increasing the DMSO concentration to 0.5% at 10 µM increased solubility allowing for concentration-response testing and 0.5% DMSO was used as vehicle control for Cmpd 7 and Cmpd 46 studies. To control for day-to-day variation in cell number and transfection, SKF-81297 was used as a positive control D1R agonist in every assay. The raw light counts per second of test compounds were normalized to the maximal SKF-81297 response (% SKF-81297 response). Data shown are from at least three independent experiments (n ≥ 3) performed in technical triplicate.

For D5R cAMP assays, HEK293 cells were transfected with 0.25 µg D5R-Stop plus 1.0 µg 22F-Glosensor plasmid and 10 µL Lipofectamine2000 (Invitrogen, Carlsbad, CA) per well of a 6-well plate. The Glosensor assay was then carried out similarly to that described for the D1R above, testing eleven-point concentration-responses (1 pM to 30 µM) of SKF-81297, A-77636, PF-1119 or Cmpd 19. 0.3% DMSO was used as vehicle control (basal) which matched the concentration of DMSO in the 30 µM compound. SKF-81297 was used as a positive control D5R agonist in every assay. The raw light counts per second of test compounds were normalized to the maximal SKF-81297 response (% SKF-81297 response). Data shown are from at least three independent experiments (n ≥ 3) performed in technical triplicate.

### β-arrestin recruitment Tango assay

The assay procedure was adapted from Kroeze et al., (Kroeze et al., 2015). Briefly, HTLA cells (HEK293 cells stably expressing a tTA-dependent luciferase reporter gene and a β-arrestin2-TEV protease fusion protein) were split at 375,000 cells per well into 6-well plates and incubated for 24 hours. The HTLA cells were then transfected using calcium phosphate transfection method. Briefly, 1.4 µg of DRD1-TANGO or DRD5-TANGO plasmid DNA (Plasmids #66268 or #66272, Addgene, Watertown, MA) and 120 µM CaCl_2_ (diluted with H_2_O from 2 mM stock) were mixed in one centrifuge tube. In a separate tube, an equal volume of 2X HEPES buffered saline (280 mM NaCl, 10 mM KCl, 1.5 mM Na_2_HPO_4_, 50 mM HEPES) was added. The contents of the tubes were then mixed together and vortexed. The mixture was added immediately, dropwise, to the HTLA cells which were returned to the incubator overnight. The transfected HTLA cells were split into a poly-L-lysine (R&D systems, Minneapolis, MN) coated 96-well white clear bottom cell culture plates (Greiner Bio-One, Monroe, MS) at 80,000 cells per well and returned to the incubator. Forty-eight hours post transfection, the HTLA cells were treated with the indicated agonist using an 11-point concentration-response (10 pmol to 30 µM) curve. Due to the long treatment time, agonists were diluted in 1X HBSS containing 1 mM ascorbic acid (100 μM in well final concentration) to reduce oxidation of compounds. After 18-20 hours of treatment, the cells were lysed using the Bright-Glo luciferase substrate, (Promega, Madison, WI) diluted 20-fold in 1X HBSS for 20 minutes at room temperature and total luminescence read using the Microbeta2 plate reader (Perkin Elmer, Waltham, MA) detecting total light counts in one second in each well. SKF-81297 was included as a positive control D1R and D5R agonist in every assay to control for cell number and transfection variation across experimental days. The raw light counts per second of test compounds were normalized to the maximal SKF-81297 response (% SKF-81297 response). The data shown are from at least three independent biological experiments (n ≥ 3) performed in technical triplicate.

### β-arrestin-GFP recruitment Total Internal Reflection Fluorescence (TIRF) microscopy

HEK293 cells were split into 6-well plates at 425,000 cells per well and incubated for 24 hours. The cells were co-transfected with 0.25 µg of HA-D1R and 0.25 µg of β-arrestin2-GFP (plasmid # 35411, Addgene, Watertown, MA) in 10 µL of lipofectamine2000 per well as directed by the manufacturer’s protocol and incubated overnight. The cells were then split into poly-L-lysine (R&D Systems, Minneapolis, MN) coated 35 mm dishes with a No. 1.5 coverslip base (MatTek Corporation, Ashland, MA) at 200,000 cells per dish. While splitting into the 35mm dishes, the cells were transferred and subsequently cultured in DMEM containing 10% dialyzed fetal bovine serum (Omega Scientific, Tarzana, CA) and 1% penicillin/streptomycin. Forty-eight hours after transfection, the cells were serum starved in 1X HBSS for 60 minutes and treated with 3 µM of the indicated agonist for 10 minutes at 37°C. The cells were fixed using 4% paraformaldehyde (Electron Microscopy Sciences, Hatfield, PA) for 20 minutes at room temperature. The dishes were washed 3 times in phosphate buffered saline (PBS) for 5 minutes each then permeabilized with 0.1% Triton X-100 (Sigma, St. Louis, MO) in PBS for 40 minutes at room temperature. The cells were again washed 3 times for 5 minutes each in PBS. The cells were blocked in 5% goat serum (Sigma, St. Louis, MO) diluted in PBS for 1 hour at room temperature. Anti-HA antibody (1:1000 dilution, catalog #3724S, Cell Signaling, Danvers, MA) diluted in 5% goat serum was incubated with the cells overnight at 4°C. The cells were then washed 3 times for 5 minutes each in PBS. Anti-rabbit Alexa594 antibody (1:250 dilution, catalog #A11012, Invitrogen, Carlsbad, CA) diluted in PBS containing 0.01% Hoechst 33342 fluorescent nuclear stain (ThermoFisher Scientific, Waltham, MA) was incubated with the cells for 60 minutes at room temperature. The cells were washed 3 times for 5 minutes in PBS. TIRF microscopy was performed using a Nikon Eclipse Ti2 dual epi fluorescence and confocal microscope outfitted with a TIRF imaging apparatus. Images were obtained using a 60x TIRF objective and dedicated laser lines for the 488 nm (β-arrestin2-GFP) or 561 nm (Alexa594 labeled HA-D1R). Image acquisition settings including laser power and exposure times were initially set using SKF-81297 treated cells (positive control) and identical settings were subsequently used for all cells imaged across treatment groups. A minimum of 21 cells (≥ 7 cells/biological experiment) were imaged for each treatment group across three independent biological experiments. Representative images were adjusted for brightness and contrast, consistent with guidelines for publication, such that all pixels were adjusted equally.

### D1R endocytosis confocal imaging

HEK293 cells were plated into 6-well plates at 400,000 cells per well. After 24 hours, the cells were transfected with 0.25 µg HA-D1R plasmid and 7.5 µL Lipofectamine2000 per well as per manufacturer’s protocol and returned to the incubator overnight. The following morning cells were plated on poly-L-lysine coated (R&D systems, Minneapolis, MN) coverslips in 24-well plates at 50,000 cells per well and returned to the incubator for 24 hours. The cells were then serum starved for 1 hour in DMEM without serum or antibiotics. After aspirating the media, ice-cold 1X HBSS with 20 mM HEPES was added gently to the cells and incubated on ice for 15 minutes. Anti-HA antibody conjugated to Alexa488 (1:200 dilution, catalog #2350S, Cell Signaling, Danvers, MA) was diluted in HEPES/HBSS and then added to the cells for 45 minutes in the dark on ice. After 3 washes in ice-cold HEPES/HBSS for 5 minutes each, 10 µM of the indicated agonist was added to the cells which were returned to 37°C for 60 minutes. The cells were fixed immediately in 4% paraformaldehyde (Electron Microscopy Sciences, Hatfield, PA) diluted in PBS for 20 minutes, followed by 3 washes in PBS for 5 minutes. The coverslips were mounted on slides using Vectorshield mounting medium with 4′,6-diamidino-2-phenylindole (DAPI, Vector Laboratories, Burlingame, CA). Cells were imaged with a 63X oil objective on a Leica True Confocal Scanner SPE and Leica Application Suite Advanced software (Leica Microsystems, Wetzlar, Germany). Z-stacks were obtained with 0.2 µm steps with a zoom factor of 1.5. Image acquisition settings including laser power and exposure times were initially set using SKF-81297 treated cells (positive control) and identical settings were subsequently used for all cells imaged across treatment groups. A minimum of 21 cells (≥ 7 cells/biological experiment) were imaged for each treatment group across three independent biological experiments. Representative images were adjusted for brightness and contrast, consistent with guidelines for publication, such that all pixels were adjusted equally.

### Cell surface enzyme-linked immunosorbent assay (ELISA)

A cell surface ELISA was used to quantify agonist-induced D1R endocytosis. HEK293 cells were plated in 6-well plates at 400,000 cells per well and incubated for 24 hours at 37°C. HEK293 cells were transfected with 0.25 µg HA-D1R plasmid and 7.5 µL Lipofectamine2000 per well as per manufacturer’s protocol and incubated overnight. The cells were then split into 96-well plates coated with poly-L-lysine at 65,000 cells per well and incubated for 24 hours. The cells were serum starved for 1 hour in DMEM without serum or antibiotics then treated with 10 µM of the indicated agonist for 0-120 minutes and fixed with ice-cold 4% paraformaldehyde in PBS for 20 minutes at room temperature. Cells were washed 3 times with PBS for 5 minutes each, followed by blocking for 1 hour in 5% bovine serum albumin (Sigma, St. Louis, MO) diluted in PBS at room temperature. The plates were incubated overnight at 4°C with rabbit anti-HA antibody (1:1000, catalog #3724S, Cell Signaling, Danvers, MA) diluted in 5% bovine serum albumin. In the morning, the cells were washed 3 times with PBS for 5 minutes. Anti-rabbit horse radish peroxidase (HRP) conjugated antibody (1:1000 dilution, catalog #7074S, Cell Signaling, Danvers, MA) diluted in PBS containing 0.01% Hoechst 33342 (ThermoFisher Scientific, Waltham, MA) fluorescent nuclear stain was added to the cells for 1 hour at room temperature. After 3 washes with PBS for 5 minutes each, Hoechst staining a was read using an excitation of 392 nm and emission of 440 nm on a Synergy H4 plate reader (Biotek, Winooski, VT). Hoechst staining was used as a measure of total cell density/well. The cells were then incubated in 3’3,5’5-tetramethylbenzidine (TMB) HRP liquid substrate (Sigma, St. Louis, MO) in the dark at room temperature for 30 minutes and then absorbance was measured at 370 nm on the Synergy H4 plate reader. The cell surface ELISA was analyzed as follows. The background absorbance was determined by measuring non-transfected cells and the mean background of non-transfected cells was subtracted from each well. Each well was then normalized to the Hoechst signal from that same well to account for cell number variation by dividing the background subtracted absorbance by the Hoechst signal. Following this, all wells were normalized to the untreated control cells by dividing each well by the mean of the Hoechst normalized untreated control cells. The control cells were defined as the untreated cells (0-minute treatment) when there was no agonist-induced D1R endocytosis. Multiplying the normalized wells by 100 converted the values to the percentage of the surface HA-D1R where 100% was the 0-minute time point (control cells) when there was no agonist-induced receptor endocytosis and decreased percentages indicate D1R endocytosis. These values were subtracted from 100% to obtain the percent loss cell surface HA-D1R (i.e. after 0-minute treatment there was 0% loss of cell surface HA-D1R). Area under the curve analysis was applied to determine total HA-D1R endocytosis as described in the statistical methods section. The data shown are from three independent biological experiments (n = 3) performed in technical triplicate.

### Saturation Binding

Stably transfected D1R or transiently transfected HTLA cells were washed with ice cold PBS. Membranes were collected by scraping in 50 mM HEPES, 50 mM NaCl, 5 mM MgCl_2_, and 0.5 mM EDTA. Harvested membranes were centrifuged three times at 4000 x g at 4 °C for 20 minutes and were frozen in -80°C for binding assays. Saturation binding isotherms were performed in 96-well plates using similar methods as previously reported (Land et al., 2019). For saturation binding assays, 0.2 to 16 nM of [^3^H]SCH23390 (83.2 Ci/mmol, Lot #2551102 PerkinElmer, Waltham, MA) was used to obtain affinity (K_d_) and receptor concentration (B_max_) values. Non-specific binding was determined in the presence of 10 µM of SCH23390 (Sigma Aldrich, St. Louis MO). The reaction mixtures were incubated with ∼ 10 µg of membrane preparations at room temperature for 90 minutes on a plate shaker in the dark to reach equilibrium, and then passed rapidly through a printed filtermat soaked in 0.5% polyethylenimine using a FilterMate Harvester (PerkinElmer, Waltham, MA). The printed filtermat containing bound [^3^H]SCH23390 was microwaved for one minute to dry, then a MeltiLex sheet was melted onto the printed filtermat via a hotplate. The contents were sealed and counted for scintillation using a MicroBeta2 (PerkinElmer, Waltham, MA). Direct radioligand concentrations were measured by pipetting into 1 mL of Optiphase Supramix (PerkinElmer, Waltham, MA) and measured on a Tri-Carb 2910TR liquid scintillation analyzer (PerkinElmer, Waltham, MA). Protein concentrations were determined using the bicinchoninic acid (BCA) protein assay kit (Thermo Scientific, Waltham, MA) by measuring absorbance values (562 nm) on a H4 synergy reader (Biotek, Winooski, VT). The data shown are from three independent biological experiments (n = 3) performed in technical triplicates.

### Statistical analysis

Concentration-response analyses were conducted using Graphpad Prism8 (GraphPad Software, San Diego, CA). Concentration-response curves were generated using least squares nonlinear regression curve fitting and the “dose-response – stimulation” four parameter fit analysis in Prism. Raw data results were normalized to the maximal SKF-81297 (100%) response in each assay and are presented as % of SKF-81297. Saturation binding isotherms are presented as B_max_ and K_d_ values, as computed by GraphPad Prism using specific binding with Hill slope nonlinear regression curve-fitting algorithm.

For bias factor calculations, mean efficacy (E_max_) and potency (EC_50_) were obtained from at least three independent concentration-responses for all agonists in cAMP and β-arrestin recruitment assays. We used the bias factor calculation and formulas from Kenakin (2017). Briefly, E_max_ and EC_50_ values for each agonist were entered into the log(E_max_/EC_50_) equation for both cAMP or Tango β-arrestin recruitment assays and then subtracted from the log(E_max_/EC_50_) for the reference full D1R agonist SKF-81297, to obtain a Δlog(E_max_/EC_50_). To obtain a ΔΔlog(E_max_/EC_50_), the Δlog(E_max_/EC_50_)_β-arrestin_ was subtracted from Δlog(E_max_/EC_50_)_cAMP_. The inverse log of ΔΔlog(E_max_/EC_50_) is the G protein bias factor. The β-arrestin bias factor is obtained by subtracting the Δlog(E_max_/EC_50_)_cAMP_ from Δlog(E_max_/EC_50_)_β-arrestin_ to calculate the ΔΔlog(E_max_/EC_50_). The β-arrestin bias factor is the inverse of ΔΔlog(E_max_/EC_50_). The Kenakin (2017) bias factor calculation method requires an agonist to have an E_max_ ˃ 35% compared to the reference agonist. Thus, a bias factor could not be calculated (denoted N.C.) for the majority of agonists tested since many agonists were extremely G protein biased with little or no β-arrestin recruitment activity.

For the cell surface ELISA, area under the curve analysis was conducted for each biological replicate using GraphPad Prism 8 to determine total HA-D1R endocytosis. The area under the curve analysis included all time points from 0-120 minutes. The resulting area under the curve analysis was plotted in a bar graph resulting from the three biological replicates (n = 3). The total HA-D1R endocytosis was analyzed in GraphPad Prism 8 using a one-way ANOVA with Bonferroni’s multiple comparisons test comparing all agonists to SKF-81297.

A Spearman’s correlation was conducted on the β-arrestin recruitment E_max_ (x) and Total HA-D1R endocytosis area under the curve (y) in GraphPad Prism 8. The non-parametric Spearman’s correlation analysis was conducted because the data was normalized preventing the use of the parametric Pearson’s correlation. Spearmen’s r and the p value are reported in the scatterplot. In addition, a linear regression line was plotted on the graph and R^2^ goodness of fit calculated using GraphPad Prism 8 to assess the linear relationship between β-arrestin recruitment (x) and Total HA-D1R endocytosis (y).

## Results

### Functional selectivity of D1R catechol agonists

Biased signaling or functional selectivity has been reported at receptors and several catechol D1R agonists were found to exhibit G protein biased signaling activity at the D1R (Conroy et al., 2015; Yang et al., 2018). We further examined select catechol D1R agonists to quantitatively assess functional selectivity and calculate bias factors where possible. First, several benzazepine-based catechol agonists were tested for D1R-G protein or D1R-β-arrestin activity in concentration-response assays. D1R-G protein activity via cAMP production was measured using the Glosensor assay in HEK293 cells stably expressing the D1R (D1-HEK). D1R β-arrestin recruitment was measured using the Tango assay also using a HEK293 cell line (HTLA cells). Therefore, a similar cellular background was used to characterize and compare D1R agonist pharmacology in the two pathways. Notably, the Tango β-arrestin assay provides a gene-reporter based readout of β-arrestin recruitment, requiring ligand incubation for approximately 18 hours (Kroeze et al., 2015). We initially validated our assays testing the endogenous agonist dopamine, which was potent and efficacious for D1R in the cAMP assay but did not saturate in concentration-responses using the Tango β-arrestin assay (**supplemental Fig. 1, Table 1**). Poor dopamine activity in the Tango assay is likely due the instability of the ligand and its known oxidation in aqueous solution over 18 hours (Sulzer and Zecca, 2000). Therefore, all efficacy measurements in concentration-response assays were normalized to the full agonist SKF-81297, which showed saturating responses in both cAMP and β-arrestin assays. All potency (EC_50_) and efficacy (E_max_) values for cAMP and β-arrestin recruitment Tango assays as well as the calculated bias factors are reported in **Table 1**. SKF-81297 showed potent full agonism for cAMP production (EC_50_ = 3.3 nM, E_max_ = 100%,) as well as full agonism for β-arrestin recruitment (EC_50_ = 280 nM, E_max_ = 100%, **Fig. 1A**). The catechol agonists SKF-38393 (EC_50_ = 110 nM, E_max_ = 83%, **Fig. 1B**) and SKF-77434 (EC_50_ = 140 nM, E_max_ = 44%, **Fig. 1C**) displayed high and low D1R partial agonism for cAMP production, respectively. Notably, SKF-38393 and SKF-77434 displayed no activity in the D1R-β-arrestin recruitment assay, in stark contrast to SKF-81297 (**Fig. 1B and C**). These results confirmed that catechol agonists SKF-38393 and SKF-77434 are G protein biased D1R agonists, consistent with a previous report (Conroy et al., 2015). Notably, the Kenakin 2017 bias factor method cannot calculate bias factors for strongly biased agonists because a minimum efficacy of 35% is required in all assays (Kenakin, 2017). Therefore, bias factor calculations were not possible for the purely G protein biased agonists, SKF-38393 and SKF-77434, consistent with their lack of activity for β-arrestin. We also examined the catechol agonist A-77636, which exhibited potent full agonism for D1R cAMP signaling (EC_50_ = 3.0 nM, E_max_ = 99%, **Fig. 1D**). Remarkably, A-77636 was more potent than SKF-81297 and unexpectedly a super-agonist for β-arrestin recruitment (EC_50_ = 34 nM, E_max_ = 130%, **Fig. 1D**). Based on this increased potency relative to SKF-81297, A-77636 had a β-arrestin bias factor of 10 and a G protein bias factor of 0.1, indicating A-77636 has β-arrestin biased activity (**Table 1**). A-77636 is a highly potent super agonist for D1R-β-arrestin activity.

**Figure 1.**
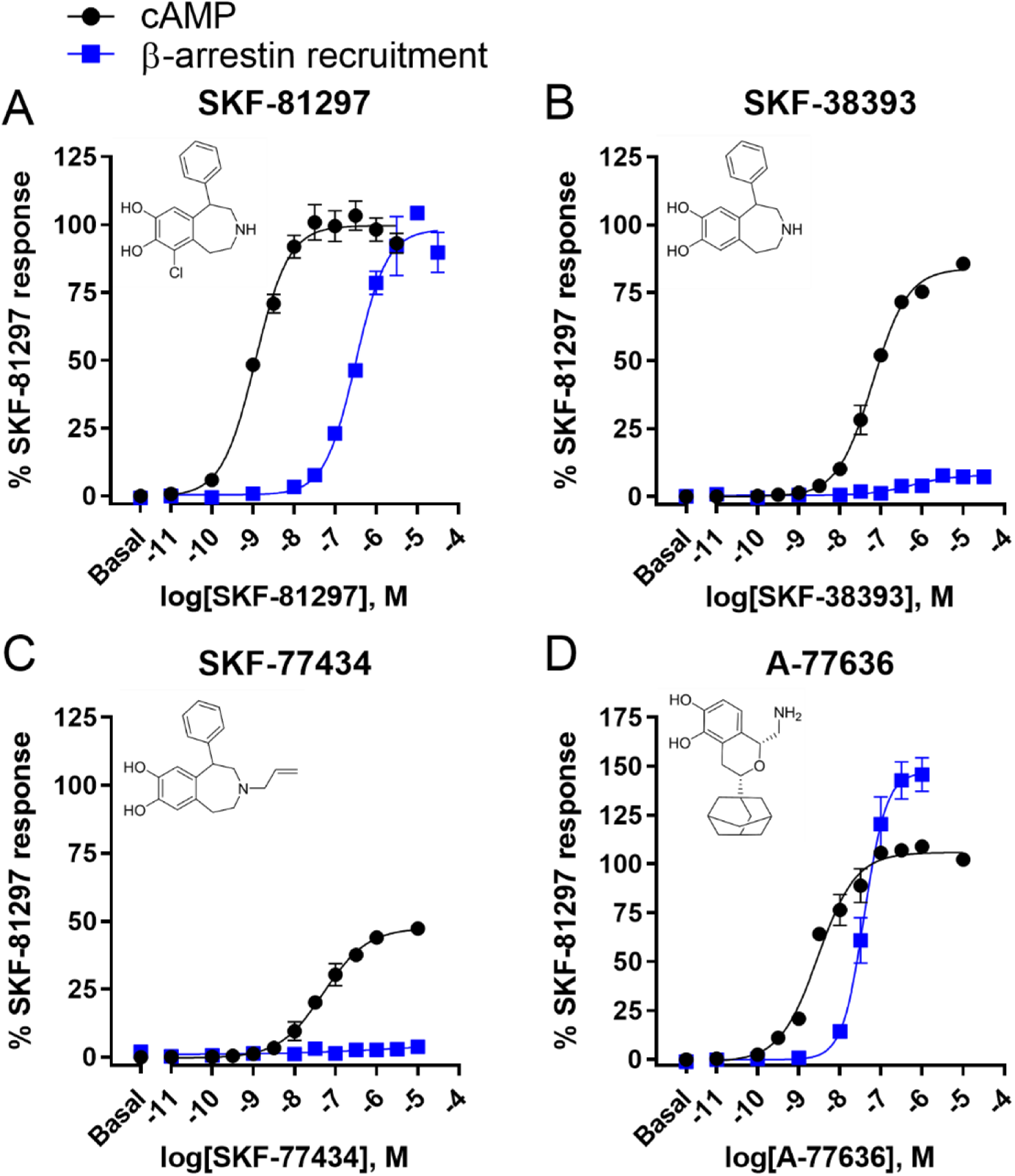
Concentration responses of catechol D1R agonists in cAMP and β-arrestin recruitment assays. D1R-mediated cAMP accumulation (black) was measured using the Glosensor assay in HEK293 cells stably expressing the D1R and β-arrestin recruitment (blue) was measured with the Tango assay as described in methods. Agonist structures are shown in upper left corner of each panel. **A)** SKF-81297 is a full agonist for both G protein/cAMP (black) and β-arrestin recruitment (blue). All agonists were normalized to SKF-81297. **B)** SKF-38393 is a high partial agonist for G protein/cAMP but does not recruit β-arrestin. **C)** SKF-77434 is a low partial agonist for G protein/cAMP activity compared to SKF-81297 but has no activity in the β-arrestin recruitment assay. **D)** A-77636 is a full agonist for G protein/cAMP signaling but, interestingly, was a super agonist for β-arrestin recruitment. Representative plots shown from at least three independent experiments performed in technical triplicate. Light counts per second (bioluminescence) from both assays was normalized to 100% SKF-81297 response. EC_50_ and E_max_ values were obtained by non-linear regression analysis for all experiments and average values are summarized in Table 1.

**Table 1.** Pharmacological activities of catechol and non-catechol agonists in D1R-Gs cAMP or D1R-β-arrestin recruitment assays and calculated bias factors.

| Compound | Gs-cAMP | | β-arrestin recruitment | | β-arrestin Bias Factor | Gs Bias Factor |
| --- | --- | --- | --- | --- | --- | --- |
|  | EC<sub>50</sub> (nM) | E<sub>max</sub> (% SKF-81297) | EC<sub>50</sub> (nM) | E<sub>max</sub> (% SKF-81297) |  |  |
| SKF-81297 | 3.3 ± 2 | 100 ± 3 | 280 ± 47 | 100 ± 3 | 1 | 1 |
| SKF-38393 | 110 ± 48 | 83 ± 4 | NA | NA | NC | NC |
| SKF-77434 | 140 ± 34 | 44 ± 2 | NA | NA | NC | NC |
| A-77636 | 3.0 ± 1 | 99 ± 2 | 34 ± 6 | 130 ± 14 | 10 | 0.1 |
| PF-2334 | 12 ± 5 | 94 ± 9 | 460 ± 67 | 70 ± 10 | 1.6 | 0.63 |
| PF-1119 | 57 ± 15 | 72 ± 8 | NA | NA | NC | NC |
| PF-6142 | 22 ± 8 | 88 ± 12 | 440 ± 89 | 22 ± 1 | NC | NC |
| Cmpd 7 | 400 ± 27 | 81 ± 8 | NA | NA | NC | NC |
| Cmpd 41 | 180 ± 67 | 71 ± 2 | NA | NA | NC | NC |
| Cmpd 46 | 340 ± 25 | 78 ± 5 | NA | NA | NC | NC |
| Cmpd 19 | 4.6 ± 3 | 91 ± 3 | 120 ± 55 | 110 ± 10 | 4.1 | 0.24 |
| DA | 38 ± 13 | 93 ± 5 | NT | NT | NT | NT |

Mean ± SD, n = 3-13, NA = no activity, NC = not calculable, NT = not tested due to assay limitations.

### Functional selectivity of D1R non-catechol agonists

To further investigate D1R agonist functional selectivity, we next tested the non-catechol D1R agonists that are structurally distinct from previously describe catechol agonists. We began with retesting three selective agonists described as G protein biased by Pfizer (Davoren et al., 2018; Gray et al., 2018). In that work, PF-1119, PF-2334, and PF-6142 increased cAMP production in neurons but did not recruit β-arrestin in U2OS cells (Gray et al., 2018). These studies suggested that the non-catechol agonists displayed G protein biased activity; however, in those studies, different cellular backgrounds were used and β-arrestin recruitment assay was not tested in a concentration-responsive fashion. Therefore, to quantitatively compare PF-1119, PF-2334, and PF-6142, we resynthesized and retested these compounds in rigorous concentration-response studies using the same cell background (HEK293). PF-2334 showed potent full agonism for D1R cAMP production (EC_50_ = 12 nM, E_max_ = 94%) but had weaker potency and partial agonist activity for β-arrestin recruitment (EC_50_ = 460 nM, E_max_ = 70%, **Fig. 2A**, **Table 1**). PF-1119 (EC_50_ = 57 nM, E_max_ = 72%) and PF-6142 (EC_50_ = 22 nM, E_max_ = 88%) displayed high partial agonism for D1R cAMP production compared to SKF-81297 (**Fig. 2B and C**). However, no activity was observed in the β-arrestin recruitment assay for PF-1119 (**Fig. 2B**) while PF-6142 had very low partial activity (EC_50_ = 440 nM, E_max_ = 22%, **Fig. 2C**). These studies further confirmed that these ligands are G protein biased with PF-1119 being purely G protein biased.

**Figure 2.**
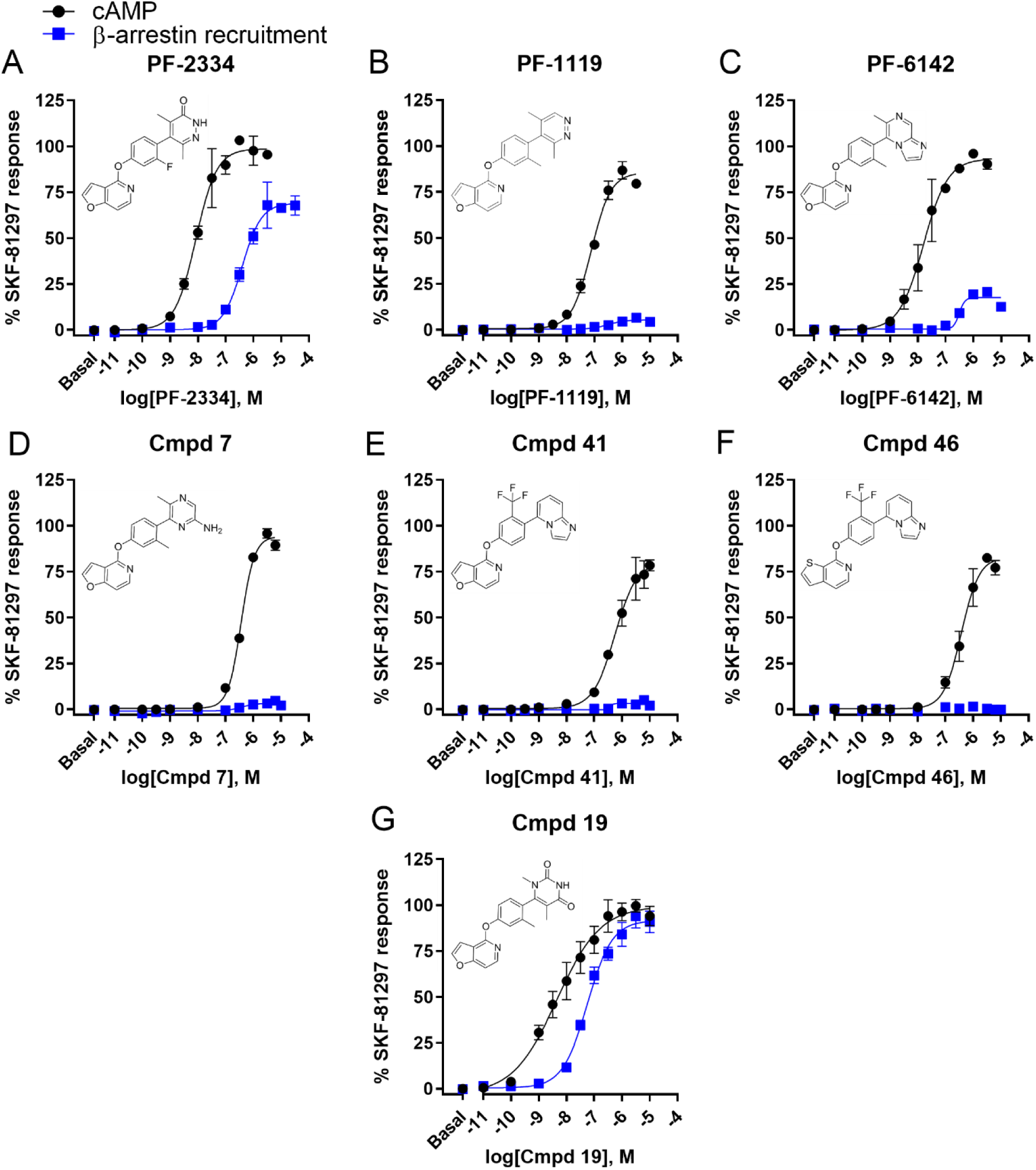
Non-catechol D1R agonist-mediated concentration responses for cAMP accumulation and β-arrestin recruitment. The Glosensor assay was used to measure cAMP accumulation in HEK293 cells stably expressing the D1R (black) while the Tango assay measured β-arrestin recruitment (blue). All responses in both assays were normalized to the full agonist, SKF-81297. Agonist structures are displayed in the upper left corner of the corresponding panel. **A)** PF-2334 is a full agonist for G protein stimulated cAMP accumulation but a partial agonist for β-arrestin recruitment. **B)** PF-1119 is a partial agonist for cAMP production and has no activity in the β-arrestin recruitment assay. **C)** PF-6142 is a partial agonist in the G protein/cAMP pathway but had very low partial agonist activity in the β-arrestin recruitment assay. **D)** Cmpd 7, **E)** Cmpd 41, and **F)** Cmpd 46 are analogs of PF-6142 from Martini et al., 2019 and are all partial agonists for D1R-mediated cAMP accumulation and do not have activity in the β-arrestin recruitment assay. **G)** Cmpd 19 is an analog of PF-2334 described in Wang et al., 2019. Cmpd 19 is a full agonist in both D1R-mediated cAMP accumulation and β-arrestin recruitment. Representative plots shown from at least three independent experiments performed in technical triplicate. Light counts per second (bioluminescence) from both assays was normalized to 100% SKF-81297 response. EC_50_ and E_max_ values were obtained by non-linear regression analysis for all experiments and average values are summarized in Table 1.

Cmpd 7, Cmpd 41, and Cmpd 46 are non-catechol analogs of PF-6142 and are suggested to have biased or unbiased activity in a recent report (Martini et al., 2019a). To further validate their D1R activity, we resynthesized and tested Cmpd 7, Cmpd 41, and Cmpd 46 here using the cAMP Glosensor and β-arrestin Tango assays. Cmpd 7 (EC_50_ = 400 nM, E_max_ = 81%) and Cmpd 46 (EC_50_ = 340 nM, E_max_ = 78%) displayed moderate potency with high partial agonism for D1R cAMP production (**Fig. 2D and F**). Cmpd 41 showed improved potency (EC_50_ = 180 nM) but had similar partial agonist efficacy (E_max_ = 71%) for D1R cAMP production (**Fig. 2E**, **Table 1**). Interestingly, Cmpd 7, Cmpd 41, and Cmpd 46 displayed no activity in the β-arrestin recruitment assay (**Fig. 2D-F**) and biased factors could not be calculated, consistent with these non-catechols being G protein biased agonists, and results were generally similar to that reported by Martini et al. (2019a). Notably, these agonists had greatly reduced potency (∼10-fold) in the D1R cAMP assay relative to their parent compound PF-6142 (EC_50_ = 22nM).

Our group recently resynthesized and characterized an analog of PF-2334, Cmpd 19 which was a potent D1R full non-catechol agonist (Coe JW et al., 2017; Coe JW et al., 2014). Here we tested Cmpd 19 using the D1R stable HEK293 cell line for cAMP production and β-arrestin recruitment. Cmpd 19 displayed high potency, high partial agonism for D1R cAMP production (EC_50_ = 4.6 nM, E_max_ = 91%, **Fig. 2G**, **Table 1**). Remarkably, Cmpd 19 also potently recruited β-arrestin but with superior efficacy to SKF-81297 (EC_50_ = 120 nM, E_max_ = 110%, **Fig. 2G**, **Table 1**). These results unexpectedly showed that Cmpd 19 has full D1R β-arrestin recruitment activity compared to all other non-catechol agonists studied here. Bias factor calculations suggested Cmpd 19 is β-arrestin biased, due to increased β-arrestin potency and efficacy when compared to SKF-81297 (β-arrestin bias factor = 4.1, **Table 1**). However, we suggest Cmpd 19 is best described as a balanced agonist, due to the ligand’s high potency and full efficacy at both D1R signaling pathways. Taken together, these results further characterize non-catechol D1R agonists, which generally are G protein biased with the exception of the balanced agonist Cmpd 19. While it could be tempting to attribute G protein bias of the previously described molecules to differences in receptor expression between cell lines used in the respective assays, the cell line in the Tango assay actually displayed ∼10-fold higher receptor levels (B_max_ = 14021 fmol/mg) than the stable line used in the cAMP assays (B_max_ = 1924 fmol/mg) (**Supplemental Fig. 2, Supplemental Table 1**).

### Functional selectivity is conserved at the dopamine D5 receptor

To date, all D1R selective agonists also possess affinity and activity at the highly homologous dopamine D5 receptor (D5R) (Felsing et al., 2019; Gray et al., 2018). To examine if agonist functional selectivity is conserved at the D5R, we tested exemplar agonists for D5R cAMP production and β-arrestin recruitment, which to our knowledge has not been examined for non-catechol agonists or A-77636. SKF-81297, A-77636, PF-1119, and Cmpd 19 were selected for testing due to the range in functional selectivity observed at the D1R. SKF-81297 showed highly potent, full agonism for D5R cAMP production (EC_50_ = 0.45 nM, E_max_ = 100%,) and for D5R β-arrestin recruitment (EC_50_ = 55 nM, E_max_ = 100%, **Fig. 3A**, **Table 2**). A-77636 displayed potent, full agonism for both D5R cAMP production (EC_50_ = 1.1 nM, E_max_ = 103%) and D5R β-arrestin recruitment (EC_50_ = 3.6 nM, E_max_ = 100%, **Fig. 3B**, **Table 2**). Notably, A-77636 had higher potency in the D5R β-arrestin recruitment assay than SKF-81297, resulting in a β-arrestin bias factor of 37. PF-1119 showed potent partial agonism for D5R-cAMP production (EC_50_ = 14 nM, E_max_ = 73%) but greatly reduced activity for D5R β-arrestin recruitment (EC_50_ = 180 nM, E_max_ = 16%, **Fig. 3C**, **Table 2**), consistent with PF-1119 being G protein biased at the D5R. Cmpd 19 showed full agonism for stimulating D5R cAMP production (EC_50_ = 3.5 nM, E_max_ = 98%, **Fig. 3D**, **Table 2**) but partial agonism for D5R β-arrestin recruitment (EC_50_ = 28 nM, E_max_ = 66%, **Fig. 3D**, **Table 2**). Cmpd 19 had higher potency than SKF-81297 for β-arrestin recruitment at the D5R, resulting in a bias factor of 11 (**Table 2**). Together, these results indicate that biased signaling is generally conserved between the D1R and the D5R for these selected agonists.

**Figure 3.**
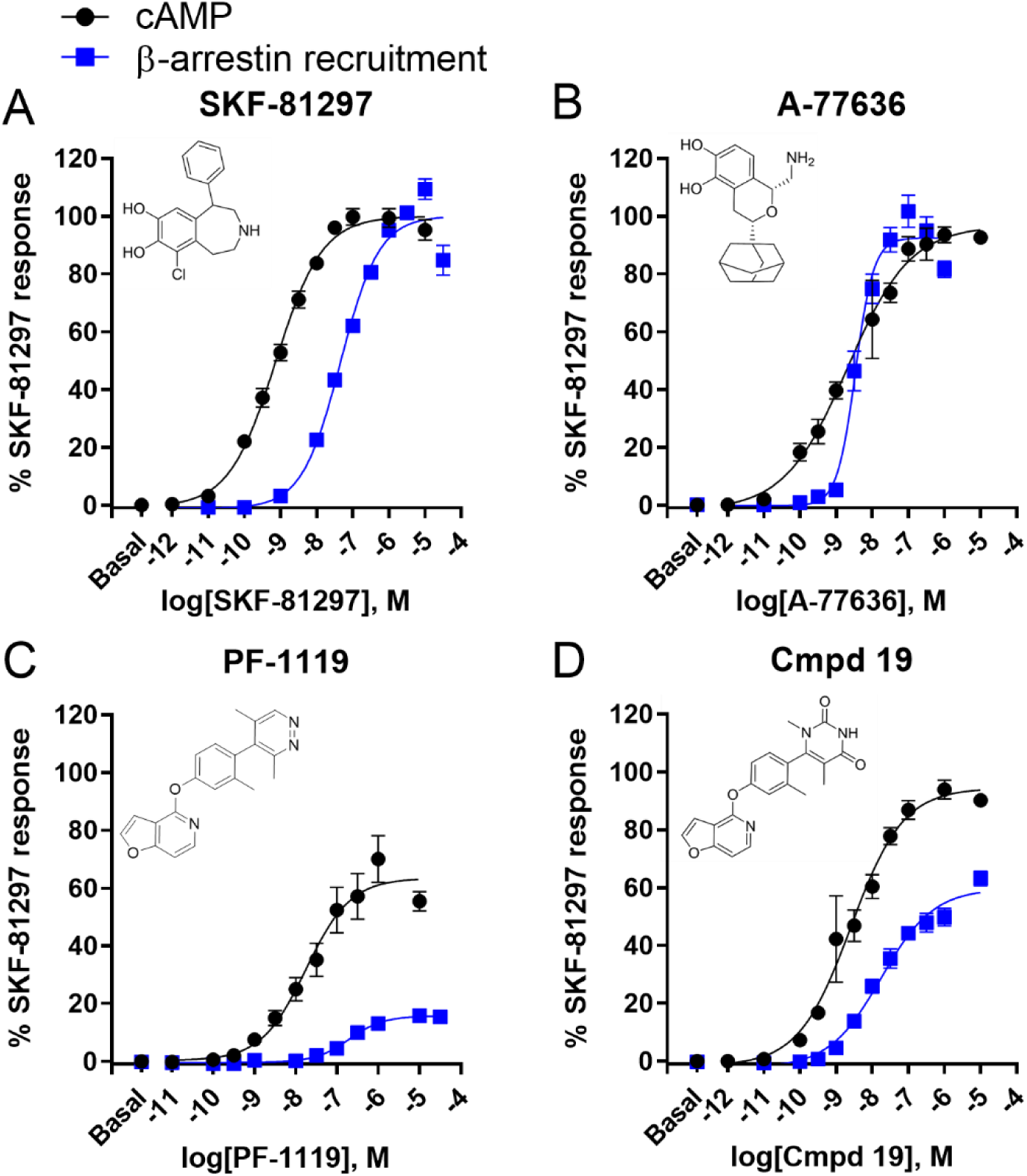
Concentration responses for <u>D5R</u>-mediated cAMP and β-arrestin recruitment. The Glosensor assay was used to measure D5R-mediated cAMP accumulation (black) in transiently transfected HEK293 cells while the Tango assay measured β-arrestin recruitment (blue). All responses in both assays were normalized to the full agonist, SKF-81297. Agonist structures are shown in the upper left corner of the corresponding panel. **A)** SKF-81297 is a full agonist in both cAMP and β-arrestin recruitment assays. **B)** A-77636 is a full agonist in both cAMP and β-arrestin recruitment assays. In the β-arrestin recruitment assay, A-77636 is more potent compared to SKF-81297. **C)** PF-1119 is a partial agonist for D5R-mediated cAMP accumulation and has very low partial agonist activity in the β-arrestin recruitment assay for the D5R. **D)** Cmpd 19 is a full agonist for D5R-mediated cAMP accumulation and is a partial agonist for β-arrestin recruitment. Representative plots shown from at least three independent experiments performed in technical triplicate. Light counts per second (bioluminescence) from both assays was normalized to 100% SKF-81297 response. EC_50_ and E_max_ values were obtained by non-linear regression analysis for all experiments and average values are summarized in Table 2.

**Table 2.** D5R EC_50_ and E_max_ values for Gs-cAMP accumulation and β-arrestin recruitment assays and corresponding β-arrestin and Gs bias factor.

| Compound | Gs-cAMP | | β-arrestin recruitment | | β-arrestin Bias Factor | Gs Bias Factor |
| --- | --- | --- | --- | --- | --- | --- |
|  | EC<sub>50</sub> (nM) | E<sub>max</sub> (% SKF-81297) | EC<sub>50</sub> (nM) | E<sub>max</sub> (% SKF-81297) |  |  |
| SKF-81297 | 0.45 ± 0.08 | 100 ± 3 | 55 ± 9 | 100 ± 3 | 1 | 1 |
| A-77636 | 1.1 ± 0.9 | 103 ± 5 | 3.6 ± 0.2 | 100 ± 6 | 37 | 0.03 |
| PF-1119 | 14 ± 4 | 73 ± 6 | 180 ± 69 | 16 ± 2 | NC | NC |
| Cmpd 19 | 3.5 ± 0.9 | 98 ± 6 | 28 ± 8 | 66 ± 9 | 11 | 0.09 |

Mean ± SD, n = 3-5, NC = not calculable.

### β-arrestin recruitment to the D1R using TIRF microscopy

To ensure the β-arrestin recruitment Tango assay did not miss the kinetic window for β-arrestin recruitment due to the length of the assay and to validate β-arrestin recruitment to the D1R, we visualized β-arrestin2-GFP translocation to the HA-tagged D1R using TIRF microscopy. TIRF microscopy uses an oblique angle of light to capture a thin slice at the plasma membrane to visualize translocation of β-arrestin2-GFP from the cytoplasm to the plasma membrane. HEK293 cells were co-transfected with β-arrestin2-GFP and HA tagged D1R (HA-D1R). The cells were treated for 10 minutes with 3 µM (lowest saturating concentration for all agonists) of the indicated agonist. The cells were fixed and immunofluorescently labeled for the HA-D1R then imaged using TIRF microscopy. Vehicle treatment did not induce β-arrestin translocation to the plasma membrane as indicated by a lack of green puncta (**Fig. 4A**). In contrast, SKF-81297 and A-77636 treatment robustly recruited β-arrestin to the plasma membrane indicated by green puncta and was colocalized with HA-D1R as seen by yellow, colocalized puncta in the merged image (**Fig. 4B and C**). As expected, dopamine also robustly recruited β-arrestin to the plasma membrane and the yellow puncta indicated colocalization with HA-D1R (**supplemental Fig. 2B**). The purely G protein biased non-catechol agonist, PF-1119, did not recruit β-arrestin to the plasma membrane (**Fig. 4D**). Cmpd 41 induced moderate β-arrestin recruitment to the plasma membrane in contrast to the lack of activity observed in the Tango assay (**Fig. 4E**). Cmpd 19 strongly induced β-arrestin recruitment to the plasma membrane and colocalized with HA-D1R (**Fig. 4F**). The catechol agonists, SKF-38393 and SKF-77434, did not recruit β-arrestin to the plasma membrane (**supplemental Fig. 2C and D**), consistent with the lack of activity in the Tango assay. PF-2334 moderately recruited β-arrestin to the plasma membrane consistent with partial β-arrestin agonism (**supplemental Fig. 3B**). The remaining non-catechol agonists did not robustly induce β-arrestin translocation aligning with the results obtained in the Tango β-arrestin assay (**supplemental Fig. 3C, D, E**). The D1R β-arrestin translocation TIRF microscopy study validated the D1R Tango β-arrestin recruitment results with one exception, Cmpd 41. Cmpd 41 displayed moderate β-arrestin recruitment in TIRF imaging but did not have activity in the Tango assay, highlighting the importance of both assays for investigating β-arrestin activity.

**Figure 4.**
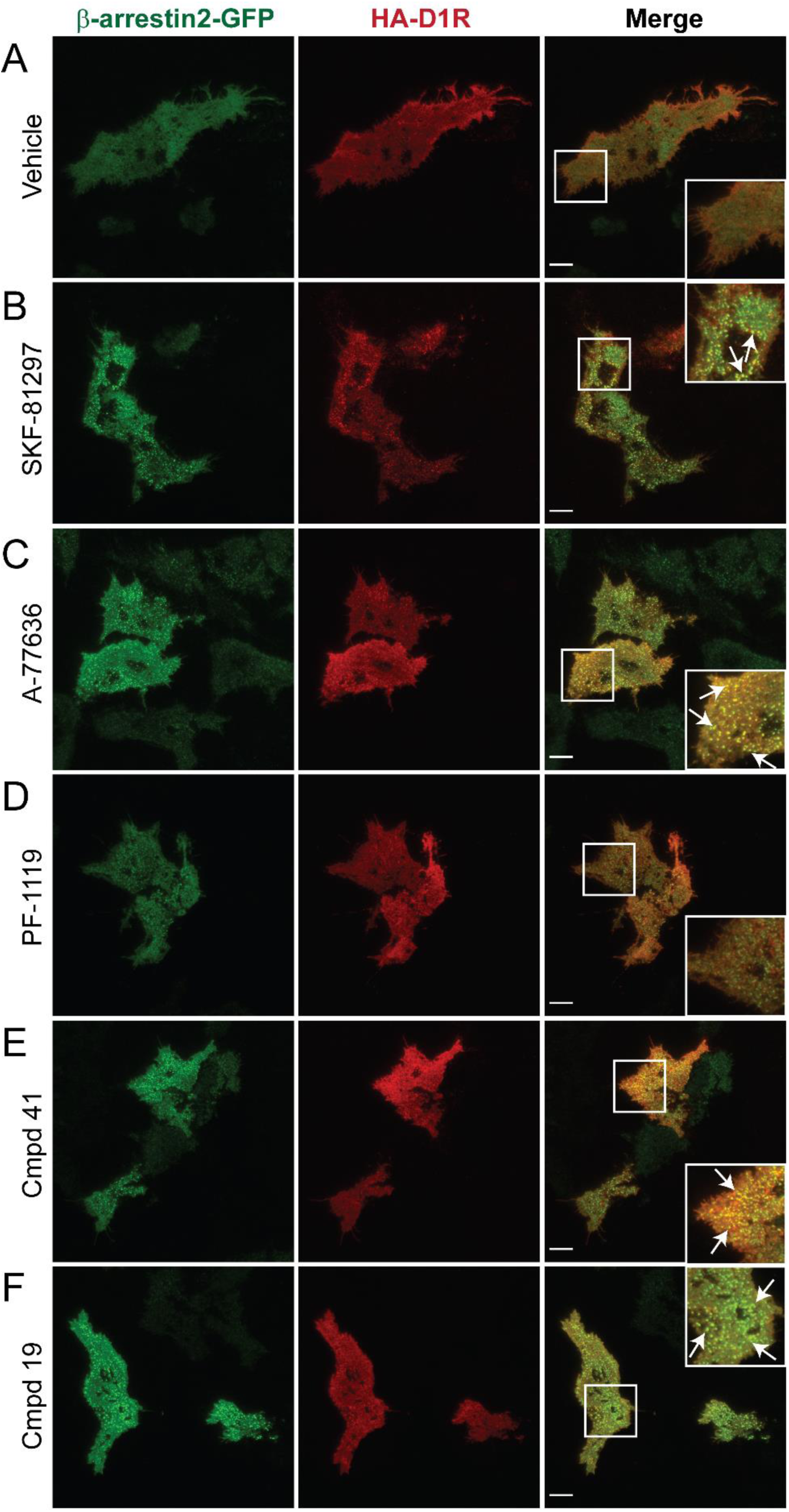
β-arrestin recruitment to the plasma membrane at a physiologically relevant time is generally consistent to the Tango assay. HEK293 cells were transfected with HA-D1R and β-arrestin2-GFP. The cells were then treated with 3 µM of the indicated agonist and fixed after 10 minutes. The D1R was then detected using antibodies against the HA tag and visualized with alexa594 secondary antibodies. TIRF microscopy was used to image a thin slice of the cells at the plasma membrane. Punctate GFP signal indicates β-arrestin recruitment to the plasma membrane. **A)** Vehicle treatment did not recruit β-arrestin to the plasma membrane. **B)** SKF-81297 strongly recruited β-arrestin to the plasma membrane. β-arrestin colocalized in puncta with HA-D1R. **C)** A-77636 treatment induced β-arrestin recruitment to the plasma membrane that colocalized with the HA-D1R. **D)** PF-1119 weakly recruited β-arrestin to the plasma membrane. **E)** Cmpd 41 moderately recruited β-arrestin to the plasma membrane. **F)** Cmpd 19 strongly recruited β-arrestin to the plasma membrane and colocalized with HA-D1R. Similar results observed in >21 cells across three independent experiments. White boxes indicate the area that is enlarged in the corner of merged images. Scale bar = 10 μm.

### D1R endocytosis using confocal microscopy

Many GPCRs undergo agonist-induced endocytosis through clathrin coated pits. A key functional outcome of β-arrestin recruitment is to allow GPCRs to be organized into clathrin coated pits and endocytosed (Ferguson et al., 1996; Goodman et al., 1996; Laporte et al., 2002; Laporte et al., 2000). The functional selectivity of the catechol and non-catechol agonists tested indicate the degree of β-arrestin recruitment varies by agonist. Thus, we hypothesized that the G protein biased agonists would not induce D1R endocytosis and that increasing levels of β-arrestin recruitment would increase D1R endocytosis. To determine the effects of balanced, G protein biased or a β-arrestin super agonist on agonist-induced D1R endocytosis, we first conducted a confocal imaging-based assay using an antibody feeding endocytosis assay. In **Fig. 5A**, the diagram illustrates the antibody feeding assay where HEK293 cells are transfected with HA-D1R. After HA-D1R expression, the live cells are chilled on ice to halt basal endocytosis and surface HA-D1Rs are labeled with an anti-HA-Alexa488 conjugated antibody as described in the methods. The cells are then warmed in the presence of 10 µM of the indicated agonist for 60 minutes, then fixed and imaged. Internalized receptor/antibody appears as a punctate signal inside the cells easily distinguished from plasma membrane localized receptors (see methods). Dopamine and SKF-81297 recruit β-arrestin and induce D1R endocytosis (Conroy et al., 2015; Gray et al., 2018) and were used as positive controls to assess D1R endocytosis. Vehicle treated cells displayed primarily plasma membrane D1R with negligible intracellular puncta indicating that the vehicle did not induce D1R endocytosis (**Fig. 5B**). Dopamine and SKF-81297 induced punctate and perinuclear antibody localization indicating agonist-induced D1R endocytosis occurred (**Fig. 5C and D**). A-77636, which showed β-arrestin super agonist activity in the Tango assay (see **Table 1**), robustly endocytosed the D1R with the strongest observed punctate and perinuclear antibody signal (**Fig. 5E**). Next, we conducted antibody feeding with several non-catechol agonists to determine the effects of G protein biased or balanced signaling on possible D1R endocytosis. PF-1119 did not induce D1R endocytosis with few puncta and no perinuclear receptor localization (**Fig. 5F**). PF-2334 had minimal intracellular puncta but not to the same robust extent as dopamine or SKF-81297 (**Fig. 5G**). This was surprising because even though PF-2334 recruited β-arrestin at 70% efficacy of SKF-81297 (**Table 1**), PF-2334 weakly induced D1R endocytosis. PF-6142 displayed minimal D1R endocytosis with little to no puncta appearing inside the plasma membrane of treated cells (**Fig. 5H**). Finally, Cmpd 19, a balanced non-catechol agonist, induced robust D1R endocytosis with punctate and perinuclear receptor localization (**Fig. 5I**). Together, these results suggest the functional selectivity of the agonist extends to D1R endocytosis. Moving forward, we decided to further investigate this agonist-induced D1R endocytosis activity of the ligands using a more quantitative and orthologous endocytosis assay.

**Figure 5.**
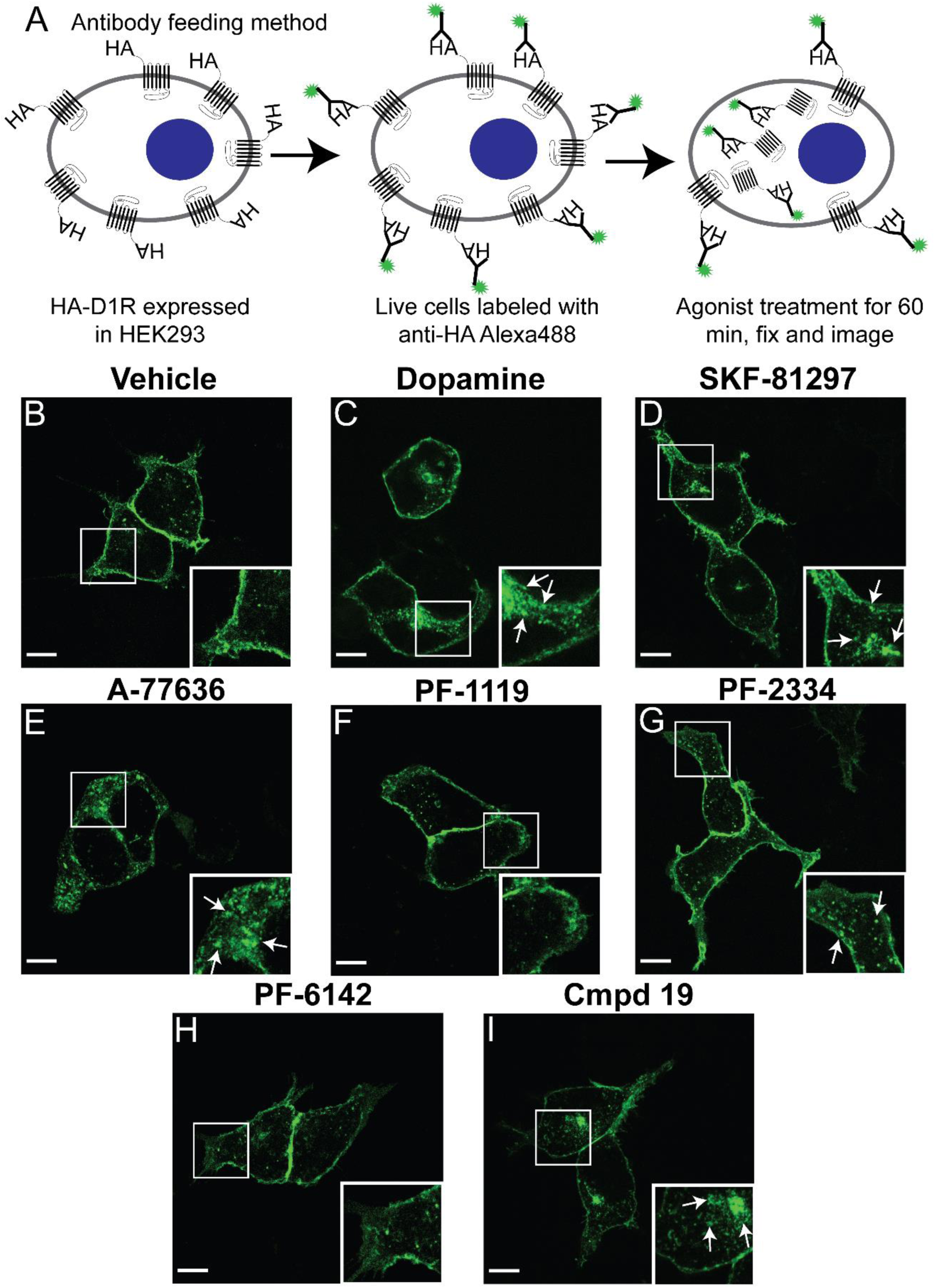
G protein biased agonists do not induce D1R endocytosis while balanced/β-arrestin super agonists do induce D1R endocytosis. **A)** A schematic representation of the Antibody Feeding assay. HEK293 cells were transfected with N-terminally HA tagged D1R. In living cells, the surface D1Rs were labeled with an anti-HA antibody conjugated to Alexa488 on ice to prevent endocytosis. The indicated agonists were added at saturating concentrations (10 μM) and the cells warmed to 37°C for 60 minutes, then fixed and imaged as described for Antibody feeding in the methods. A punctate appearance inside the plasma membrane indicates receptor/antibody endocytosis. **B)** Cells treated with vehicle for 60 minutes undergo trivial amounts of D1R endocytosis. Cells treated with the balanced catechol agonists, **C)** dopamine and **D)** SKF-81297, had punctate and perinuclear HA-D1R localization indicating dopamine and SKF-81297 strongly induced D1R endocytosis. **E)** HEK293 cells treated with A-77636 had numerous puncta and perinuclear antibody localization indicating robust D1R endocytosis. **F)** In contrast, the non-catechol agonist PF-1119 did not induce D1R endocytosis. **G)** PF-2334 induced less D1R endocytosis than SKF-81297 as seen by puncta without perinuclear signal. **H)** PF-6142 did not induce D1R endocytosis. **I)** Cmpd 19 induced robust D1R endocytosis. Representative images shown from three independent experiments. Similar results observed in >20 cells across three independent experiments. White boxes indicate the area that is enlarged in the lower right corner. White arrows indicate endocytosis. Scale bar = 10 μm.

### Quantitatively assessing D1R endocytosis through enzyme-linked immunosorbent assay

To quantitatively assess agonist-induced D1R endocytosis, a cell surface enzyme-linked immunosorbent assay (ELISA) was used to further measure activity of the various D1R ligands. The diagram in **Figure 6A** illustrates the basic cell surface ELISA process. The HA-D1R was transfected into HEK293 cells. After 48 hours, the cells were treated with a saturating concentration of agonist (10 μM) for 0-120 minutes and fixed. An ELISA was conducted to quantify the receptor remaining at the plasma membrane after agonist treatment, without permeabilizing cells to reduce detection of intracellular HA-D1R. The three catechol agonists, SKF-81297, dopamine, and A-77636 all induced D1R endocytosis (**Fig. 6B**). The maximum loss of cell surface HA-D1R induced by SKF-81297 was 33% while dopamine induced 29% loss of cell surface HA-D1R (**Fig. 6B**). Remarkably, A-77636 induced a maximum of 47% loss of cell surface HA-D1R, consistent with super agonism for β-arrestin recruitment in the Tango and TIRF assays (**Fig. 6B**). PF-1119 did not induce substantial D1R endocytosis with a maximum of 5% loss of HA-D1R from the cell surface (**Fig. 6C**). PF-6142 induced a maximum 13% loss of cell surface HA-D1R while PF-2334 induced a maximum of 16% loss of cell surface HA-D1R (**Fig. 6C**). These results align with the β-arrestin recruitment results in which PF-1119 did not recruit β-arrestin, PF-6142 was a very low partial agonist, and PF-2334 was a partial agonist for β-arrestin recruitment. Cmpd 19 induced a maximum of 28% loss of cell surface HA-D1R, similar to SKF-81297 and dopamine (**Fig. 6C**). To compare total HA-D1R endocytosis over the entire 120-minute treatment, we conducted area under the curve analysis which facilitated statistical comparisons. A-77636 induced significantly more total HA-D1R endocytosis than SKF-81297 (**Fig. 6D**, p < 0.05). The G protein biased agonists, PF-1119 and PF-6142 induced significantly less total HA-D1R endocytosis than SKF-81297 (**Fig. 6D**, p < 0.05). PF-2334 also induced significantly less endocytosis than SKF-81297 (**Fig. 6D**, p < 0.05). Dopamine and Cmpd 19 were not significantly different from SKF-81297 (**Fig. 6D**, p > 0.05). These results indicate that the balanced agonists induced similar amounts of endocytosis while the G protein biased agonists all induced less D1R endocytosis. Furthermore, A-77636, the β-arrestin super agonist, induced more D1R endocytosis than the balanced agonists. These results are consistent with the antibody feeding imaging assay.

**Figure 6.**
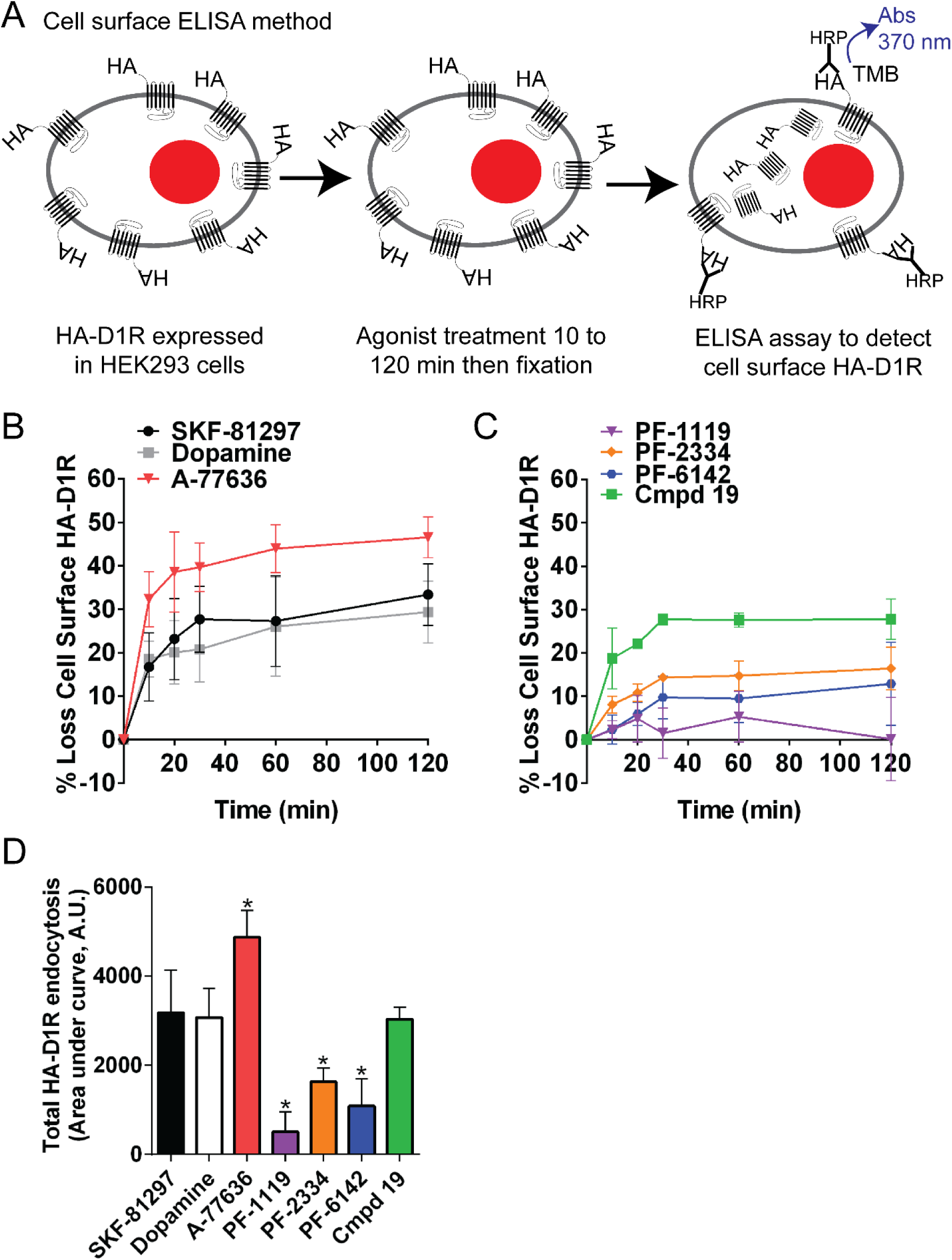
Balanced and β-arrestin super agonists induced D1R endocytosis while G protein biased agonists did not induce D1R endocytosis. **A)** HEK293 cells transfected with HA-D1R were treated with saturating concentrations (10 μM) of the indicated agonist for 0-60 minutes and fixed. Surface HA-D1Rs were detected using an ELISA assay conducted under non-permeabilizing conditions with an anti-HA antibody as described in the methods. **B)** Balanced/β-arrestin super agonists, SKF-81297, dopamine, and A-77636 induced D1R endocytosis after agonist treatment. A-77636 induced a maximum of 47% of the surface HA-D1R to be endocytosed compared to SKF-81297 and dopamine which were 33% and 29%, respectively. **C)** The non-catechol agonists induced varying levels of D1R endocytosis. PF-2334 induced a maximum of 16% HA-D1R endocytosis while PF-6142 induced a maximum of 13% HA-D1R endocytosis. PF-1119 induced the lowest HA-D1R endocytosis with a maximum of 5% HA-D1R endocytosis. Cmpd 19, the balanced non-catechol agonist, induced D1R endocytosis at similar levels to the balanced catechol agonists with a maximum D1R endocytosis of 28%. **D)** Total HA-D1R endocytosis (Area under the curve, A.U.) analysis was conducted to determine the total amount of HA-D1R endocytosis across the 120 minute treatment. A-77636 induced significantly more D1R endocytosis than SKF-81297. On the other hand, PF-1119, PF-2334, and PF-6142 induced significantly less total HA-D1R endocytosis than SKF-81297. Both dopamine and Cmpd 19 were not significantly different from SKF-81297. Data presented as Mean ± SD, n=3, *, p<0.05 vs. SKF-81297; Two-way ANOVA with Bonferroni’s multiple comparisons test.

These results support the concept that D1R agonist activity for β-arrestin recruitment and endocytosis are linked. To test this hypothesis, a Spearman’s correlation analysis was conducted between β-arrestin recruitment efficacy and total HA-D1R endocytosis (area under the curve). The degree of agonist-induced β-arrestin recruitment strongly correlates with D1R endocytosis (**Fig. 7**, r = 0.943, p = 0.008). The β-arrestin super agonist, A-77636 (upper right corner) induced more endocytosis, while the G protein biased agonists clustered in the lower left quadrant with reduced endocytosis. Thus, the higher efficacy a D1R agonist has for β-arrestin recruitment, the higher the degree of D1R endocytosis. Together, these results demonstrate D1R G protein biased agonists not only show reduced or absent β-arrestin recruitment, but that this profoundly impacts D1R endocytosis. These results further demonstrate most non-catechol D1R agonists are functionally selective with an activity profile of not only reduced β-arrestin recruitment but reduced receptor endocytosis.

**Figure 7.**
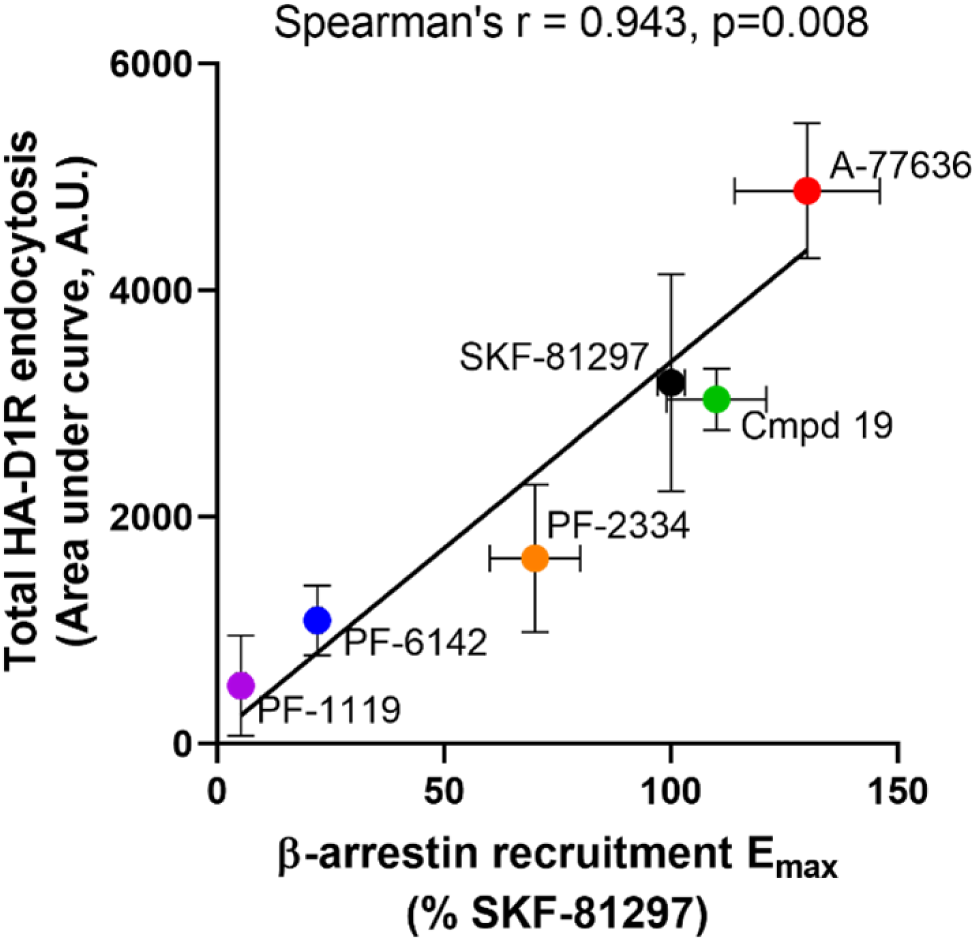
Total HA-D1R endocytosis positively correlates to β-arrestin recruitment efficacy. β-arrestin recruitment E_max_ for each agonist was plotted on the x-axis with the total HA-D1R endocytosis (area under the curve, A.U.) on the y-axis. Spearmen’s correlation coefficient indicated a strong positive correlation between β-arrestin recruitment efficacy and D1R endocytosis (r=0.829, p=0.029, Spearmen’s Correlation, one-tailed). Linear regression was performed as well to produce a line on the plot (R^2^ = 0.836, y=31.88*x = 368.0).

## Discussion

Here we demonstrate functionally selective D1R signaling and receptor trafficking by catechol and non-catechol agonists. Catechol and non-catechol D1R agonists exhibited a spectrum of signaling, ranging from highly G protein biased to balanced, despite the agonists being structurally distinct. SKF-38393 and SKF-77434 were confirmed as G protein biased, while A-77636 exhibited β-arrestin super agonism. A majority of the non-catechol agonists were G protein bias with reduced β-arrestin recruitment activity with one notable exception, Cmpd 19, which displayed balanced D1R agonism. This biased activity of profiled ligands cannot be explained by D1R expression levels in the β-arrestin cells, as they expressed ∼10-fold higher D1R versus the cells used in the cAMP assay. At the highly homologous D5R, we observed that functional selectivity is largely conserved. In an orthologous β-arrestin recruitment assay, we validated that the G protein biased agonists recruited less β-arrestin2-GFP to the D1R at the plasma membrane using TIRF microscopy. We further examined the effect of this functional selectivity on D1R endocytosis and determined G protein biased agonists induced less D1R endocytosis and that the degree of agonist-induced β-arrestin recruitment is strongly correlated to D1R endocytosis. Taken together, these results demonstrate functionally selective signaling and D1R endocytosis by both catechol and non-catechol D1R agonists.

This work defines functionally selective D1R agonists arising from structurally distinct pharmacophores. Catechol and non-catechol D1R agonists can produce a range of functional selectivity from purely G protein biased to β-arrestin super agonism. Functional selectivity is suggested to arise from an agonist’s ability to stabilize a receptor conformation that favors the activation of one transducer over another (Gray et al., 2018; Sanchez-Soto et al., 2020; Wacker et al., 2013; Wacker et al., 2017). Recently, several cryo-EM and crystal structures of the D1R were solved with the bound G protein and a variety of D1R ligands including several non-catechols (Sun et al., 2021; Teng et al., 2022; Zhuang et al., 2021). The structures of the D1R bound to the non-catechol agonists indicate the non-catechols bind within the orthosteric binding pocket although via different interactions than catechols (polar vs. hydrogen bonds, etc.) and make extended contacts with extracellular loop 2 (Gray et al., 2018; Teng et al., 2022; Xiao et al., 2021; Zhuang et al., 2021). Teng et al. (2022) suggest that β-arrestin recruitment may be determined by the distance between the ligand and TM5 in the orthosteric binding pocket. For example, in their structure, PF-06649751 (tavapadon) is further from TM5 than another non-catechol that has an additional oxygen in the structure increasing the overall length of the ligand (Teng et al., 2022; Xiao et al., 2021). PF-06649751 can recruit β-arrestin while PW0464 is highly G protein biased (Teng et al., 2022). The steric constraints put on TM5 by the length of the non-catechol ligand influences the size of the orthosteric binding pocket where a tighter pocket more efficiently recruits β-arrestin (Teng et al., 2022). Together, these structural studies suggest that the mechanism for G protein bias may have to do with the steric constraints that the ligand places on TM5. Our results indicate that catechol and non-catechol agonists can exhibit similar functional selectivity with different pharmacophores. These results further suggest that distinct chemical scaffolds are capable of stabilizing similar receptor conformations resulting in balanced or biased signaling.

These studies identified novel and unexpected activity for both catechol and non-catechol agonists. First, A-77636 is a robust β-arrestin super agonist when compared to SKF-81297. The β-arrestin super agonist activity of A-77636 was observed in two orthologous β-arrestin recruitment assays (Tango and TIRF) compared to SKF-81297. This may be due to binding kinetics, as A-77636 is known to bind to the D1R in a near irreversible manner (Lin et al., 1996) and this could potentially lock the receptor into an active conformation that facilitates β-arrestin recruitment. Interestingly, A-77636 also induced more D1R endocytosis than balanced agonists. These results are consistent with previously published results where A-77636 was shown to induce increased D1R endocytosis compared to dopamine, although the authors did not investigate β-arrestin recruitment (Lewis et al., 1998). The current study also demonstrates full β-arrestin recruitment efficacy of one non-catechol agonist, Cmpd 19. Most non-catechol D1R agonists tend to be G protein biased (Gray et al., 2018); however, our study indicates that Cmpd 19 is a balanced agonist, similar to SKF-81297. Notably, the balanced D1R activity of Cmpd 19 described here and orginally by our group (Nilson, 2020; Wang et al., 2019) has also been independently validated and the compound induced robust locomotor activity in a rodent model of Parkinson’s disease (see Cmpd 10 in (Martini et al., 2019b)). This unique balanced non-catechol D1R agonist is an invaluable tool to study the therapeutic utility of D1R biased signaling *in vivo* by comparing it with structurally similar G protein biased non-catechol agonists (e.g. PF-1119).

The D1R non-catechol agonists tested here from Martini et al. (2019a), Cmpd 7, Cmpd 41, and Cmpd 46 displayed remarkably decreased potency (∼57 - 154-fold) compared to the previously published results for G protein/cAMP production. Additionally, the authors indicated that Cmpd 41 displayed balanced agonism, which was not observed here. Cmpd 41 had no activity in the Tango β-arrestin recruitment assay; however, Cmpd 41 showed minimal activity for recruitment of β-arrestin2-GFP to the plasma membrane in TIRF microscopy. Regardless, the results here suggest that Cmpd 41 is a partial agonist for D1R β-arrestin recruitment at most. The discrepancy between the studies may be due to the different assay systems used. The present study used the Tango β-arrestin recruitment assay, which is a gene-reporter assay that requires 18-hour incubation with agonists, while Martini et al. (2019a) used a BRET assay which measures β-arrestin recruitment at much earlier time points. To ensure that the Tango β-arrestin recruitment assay did not miss the window for β-arrestin recruitment, we used an orthologous TIRF assay at similar time points to the BRET assay. Interestingly, Cmpd 41 displayed partial activity in the TIRF assay reconciling some of the discrepancies between these studies. However, we did not investigate Cmpd 41, Cmpd 7, or Cmpd 46 further due to weak potency in G protein signaling and solubility issues. The TIRF microscopy studies confirmed the results for most of the agonists tested but is not amenable to a concentration-response format highlighting the need for both assays for investigating β-arrestin recruitment at GPCRs.

The highly homologous D5R has been pharmacological indistinguishable from the D1R and functional selectivity at the D5R has not been extensively studied. Thus, we investigated if functional selectivity is conserved at the D5R for several catechol and non-catechol agonists. The select agonists (SKF-81297, A-77636, PF-1119, and Cmpd 19) displayed increased potency for both the cAMP and β-arrestin recruitment assays at the D5R compared to the D1R. More importantly, functional selectivity was conserved at the D5R for the selected agonists. Future studies investigating D5R endocytosis and desensitization are needed to characterize the impact of functional selectivity at this receptor.

Functional selectivity has renewed research efforts into multiple GPCR targets for their potential to treat a variety of disorders while displaying reduced side-effect profiles (Smith et al., 2018). An important component for functionally selective ligands is endocytosis. Since β-arrestin acts as an adaptor for clathrin-mediated endocytosis for many GPCRs (Goodman et al., 1996; Laporte et al., 2002; Laporte et al., 2000), it follows that less β-arrestin recruitment to the receptor will also decrease receptor endocytosis which, may contribute to reduced tolerance. Our results indicate that G protein biased agonists induce less D1R endocytosis compared to balanced agonists. In fact, β-arrestin recruitment efficacy strongly correlates with total D1R endocytosis. Furthermore, less receptor endocytosis suggests that fewer D1Rs are entering recycling and degradation pathways, perhaps leading to sustained D1R signaling over time since more receptors remain at the plasma membrane. Alternatively, the D1R may not be able to resensitize as rapidly since endocytosis and recycling often aid in resetting the receptor (i.e. dephosphorylation). Future studies are needed to determine the effect of G protein biased signaling and reduced D1R endocytosis on desensitization and resensitization.

In summary, our results indicate that functional selectivity at the D1R can be accomplished by structurally distinct agonists. Catechol and non-catechol agonists can exhibit biased or balanced signaling. Furthermore, functional selectivity extends to agonist-induced D1R endocytosis where G protein biased agonists do not induce D1R endocytosis while balanced and β-arrestin super agonists strongly induce receptor endocytosis. **Figure 8** summarizes these findings. Balanced agonists activate G protein signaling to increase cAMP and, in turn, recruit β-arrestin leading to D1R endocytosis (**Fig. 8A**). G protein biased agonists activate G protein signaling and increase cAMP but have reduced or even absent β-arrestin recruitment and D1R endocytosis (**Fig. 8B**). In contrast, β-arrestin super agonists activate G protein signaling and cAMP production but strongly recruit β-arrestin leading to enhanced D1R endocytosis (**Fig. 8C**). Together these results describe functionally selective D1R signaling and endocytosis by catechol and non-catechol agonists and highlight non-catechol agonists as highly valuable tools with diverse signaling and bias profiles for future studies on D1R signaling *in vitro* and *in vivo*.

**Figure 8.**
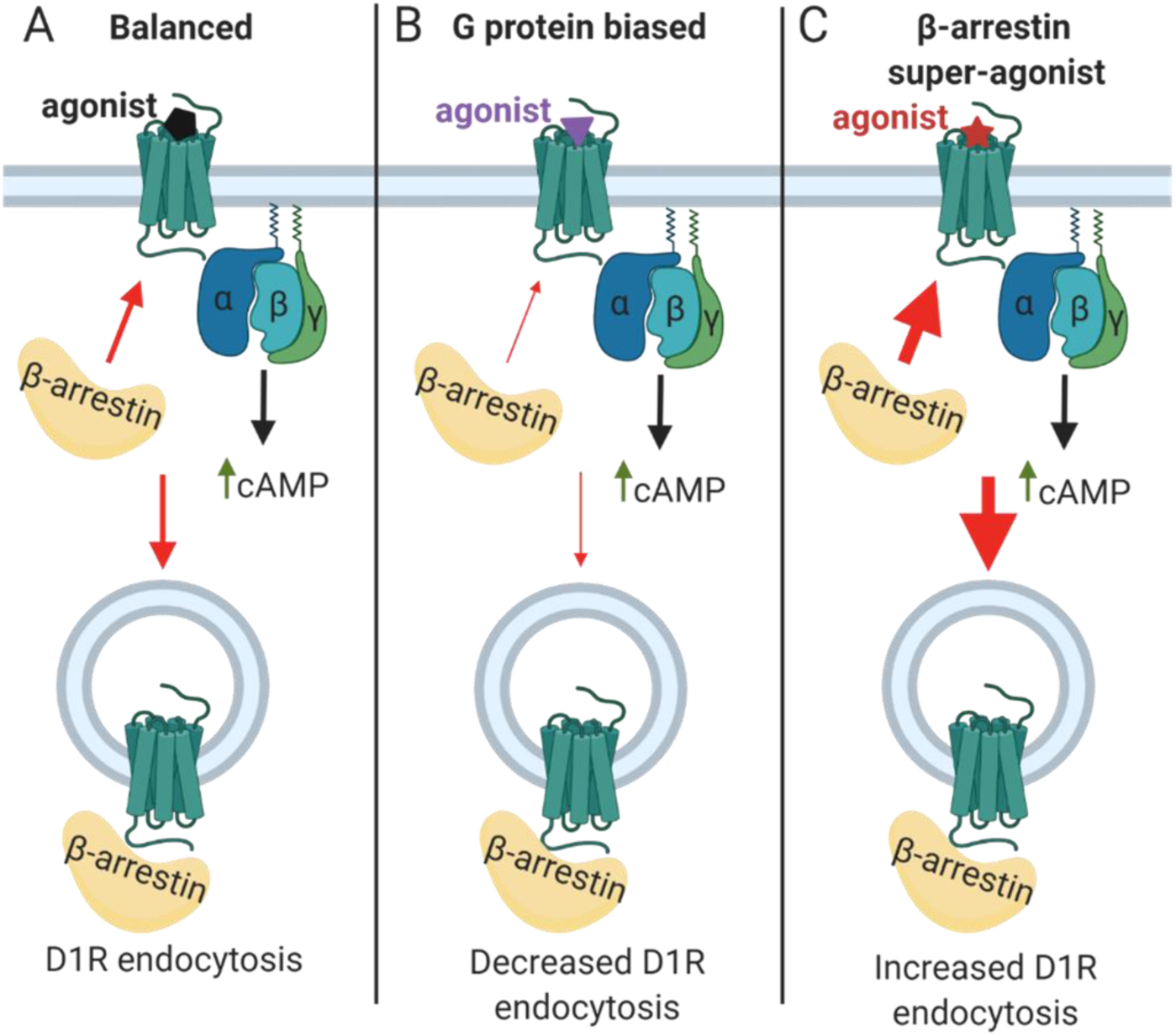
Functionally selective D1R signaling and endocytosis. The thickness of the arrow indicates the strength of the agonist to induce the indicated response. **A)** Balanced D1R agonist bind the D1R stabilizing a receptor conformation that activates the Gs/olf G protein to increase cAMP. In addition, β-arrestin can bind the D1R leading to receptor endocytosis through clathrin coated pits. **B)** G protein biased agonists stabilize a conformation that favors the activation of the G protein and cAMP production with reduced or even absent β-arrestin recruitment to the receptor. Since β-arrestin recruitment to the D1R is reduced and β-arrestin cannot act as a clathrin adaptor, D1R endocytosis is also reduced or absent. **C)** β-arrestin super agonists stabilize the receptor in a conformation that favors β-arrestin recruitment. G protein activation still occurs for D1R β-arrestin super agonists. However, β-arrestin is strongly recruited to the D1R and subsequent D1R endocytosis is enhanced.

## Abbreviations

Abs: absorbance
ANOVA: analysis of variance
A.U.: arbitrary units
B_max_: receptor concentration
Cmpd: compound
D1R: dopamine D1 receptor
D5R: dopamine D5 receptor
DMEM: Dulbecco’s modified Eagle’s medium
DMSO: dimethyl sulfoxide
EC_50_: potency
ECL: extracellular loop
E_max_: efficacy
ELISA: enzyme-linked immunosorbent assay
GFP: green fluorescent protein
GPCR: G protein-coupled receptor
HA: hemagglutinin
HBSS: Hank’s balanced salt solution
HEK293: human embryonic kidney 293
HRP: horse radish peroxidase
HTLA: HEK293 cells stably expressing tTA-dependent luciferase reporter and a β-arrestin2-TEV fusion gene
K_d_: affinity
PBS: phosphate buffered saline
TIRF: total internal reflection fluorescence
TM: transmembrane
TMB: 3’3,5’5-tetramethylbenzidine.

## Acknowledgments

We thank Dr. Bryan Roth and Dr. Wesley Kroeze (UNC Chapel Hill) for generously providing the HTLA HEK293 cells and protocols for the Tango β-arrestin assay. We also thank Dr. Casey Wright (UTMB) for generously providing access to and training for the Nikon TIRF Microscope. We also thank Sweta Raval for providing technical assistance and support in developing the β-arrestin Tango assay in our lab.

## Authorship Contributions

*Participated in research design:* Nilson, Allen

*Conducted experiments:* Nilson, Felsing, Allen

*Contributed new reagents:* Wang, Felsing, Jain, Zhou

*Performed data analysis:* Nilson, Felsing, Allen

*Wrote or contributed to the writing of the manuscript:* Nilson, Felsing, Allen

