## Supplementary figures for "Functionally selective dopamine D1 receptor endocytosis and signaling by catechol and non-catechol agonists"

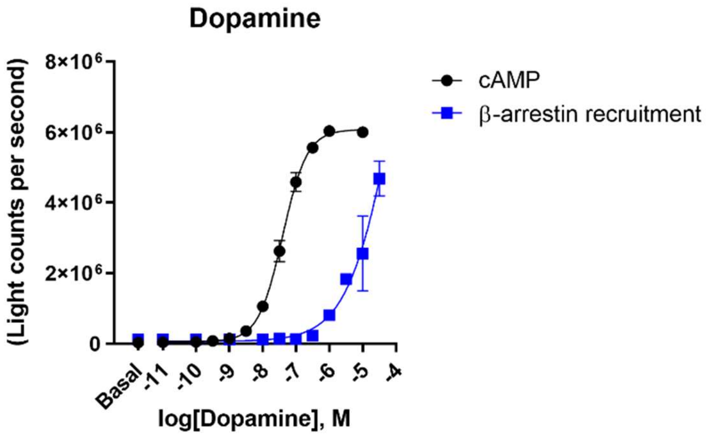


**Supplemental Figure 1.** Dopamine exhibits concentration responsive cAMP production (black) but fails to saturate in the β-arrestin recruitment assay (blue). β-arrestin recruitment was measured using the Tango assay, a gene-reporter based assay requiring 18 hours of treatment with the agonists. Dopamine is unstable and oxidizes rapidly. Even after adding 100 uM ascorbic acid to delay oxidation, β-arrestin recruitment did not saturate. This is likely due to the length of the assay being 18 hours and the degradation of dopamine. Thus, for this long assay, the endogenous agonist, dopamine cannot be used to normalize agonist responses.

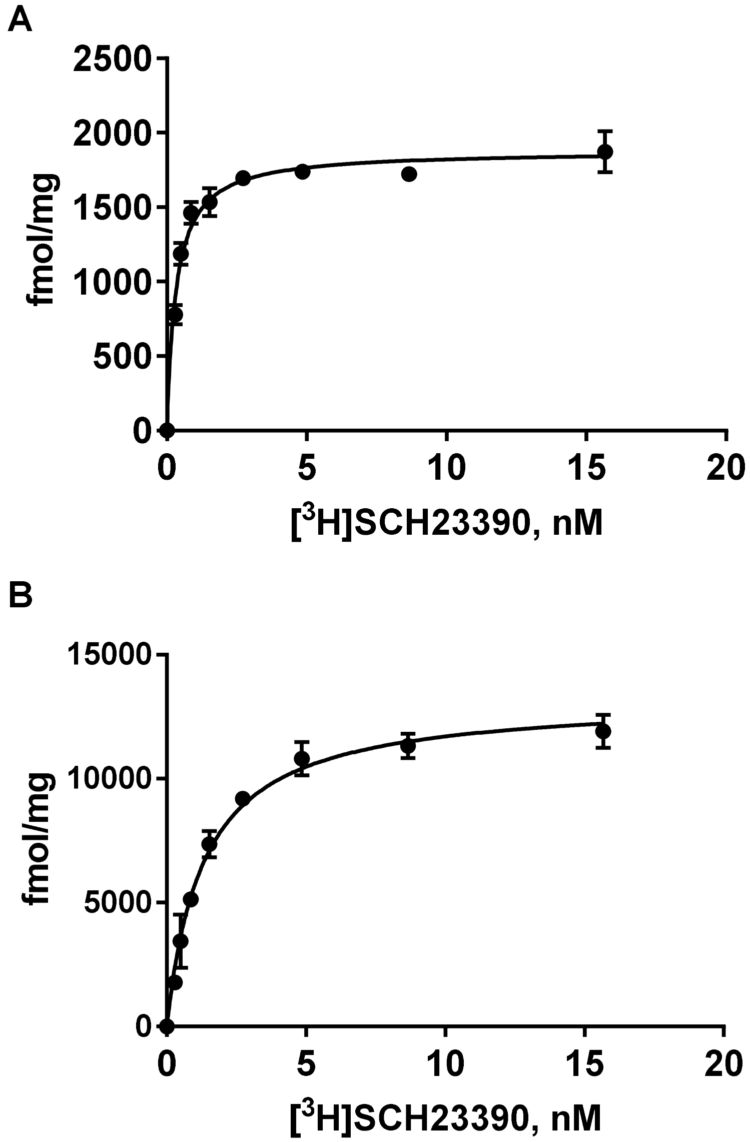


**Supplementary Figure 2. Saturation binding isotherms of cell lines used. A)** Representative saturation binding isotherm of stably transfected D1 HEK293 cell line used in cAMP assays. **B)** Representative saturation binding isotherm of transiently transfected HTLA cell line used in β-Arrestin recruitment assays. Data shown are mean ± SD using technical triplicates.

| | $K_d$ (nM) | $B_{max}$ (fmol /mg) |
|---|---|---|
| **D1 stable** | 0.33 ± 0.04 | 1924 ± 107 |
| **HTLA** | 1.48 ± 0.11 | 14021 ± 1560 |

**Supplementary Table 1. Affinity ($K_d$) and Protein Levels ($B_{max}$) of cell lines used.** Membranes (~10 ug) isolated from HEK293 (either D1 stable or transiently transfected HTLA) cells expressing the D1R were incubated with increasing concentrations of $[^3H]$SCH23390 with (non-specific binding) and without (total) the presence of cold 10 uM SCH23290. Isotherms were analyzed to determine the affinity ($K_d$) of $[^3H]$SCH23390 and total protein level ($B_{max}$) in each cell line. Data shown are the mean ± SD of biological triplicate (n=3) experiments.

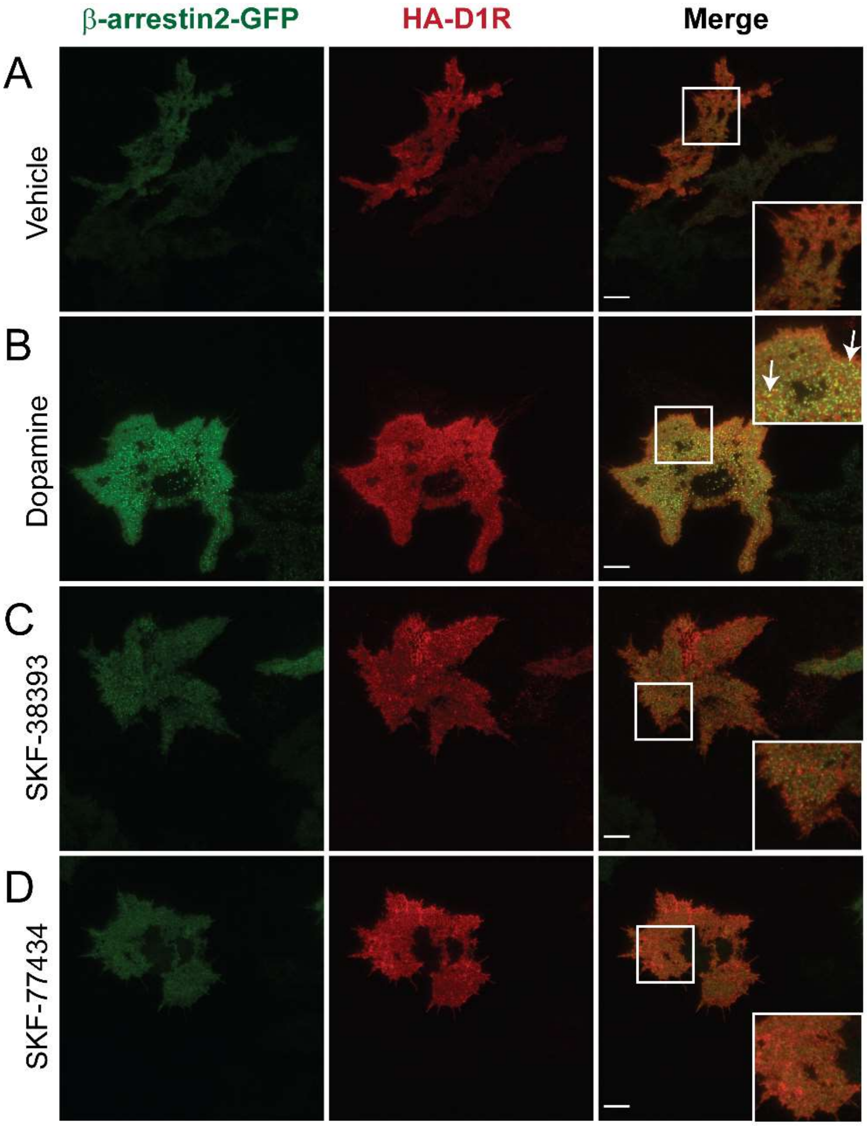


**Supplemental Figure 3. β-arrestin2-GFP recruitment to cell surface HA-D1R after treatment with catechol agonists.** HEK293 cells were transfected with HA-D1R and β-arrestin2-GFP. The cells were then treated with 3 uM of the indicated agonist and fixed after 10 minutes. The D1R was then detected using antibodies against the HA tag and visualized with alexa594 secondary antibodies. Punctate GFP signal indicates β-arrestin recruitment to the plasma membrane. TIRF microscopy was used to image a thin slice of the cells at the plasma membrane. **A)** Vehicle treatment does not recruit β-arrestin to the plasma membrane. **B)** Treating HEK293 cells with dopamine strongly induces the recruitment of β-arrestin to HA-D1Rs at the plasma membrane. β-arrestin and HA-D1R colocalize in puncta at the plasma membrane. **C)** SKF-38393 treatment did not recruit β-arrestin to the plasma membrane. **D)** Treatment with SKF-77434 did not induce the recruitment of β-arrestin to the plasma membrane. Similar results observed in >21 cells across three independent experiments. White boxes indicate the area that is enlarged in the corner of merged images. Scale bar = 10 μm.

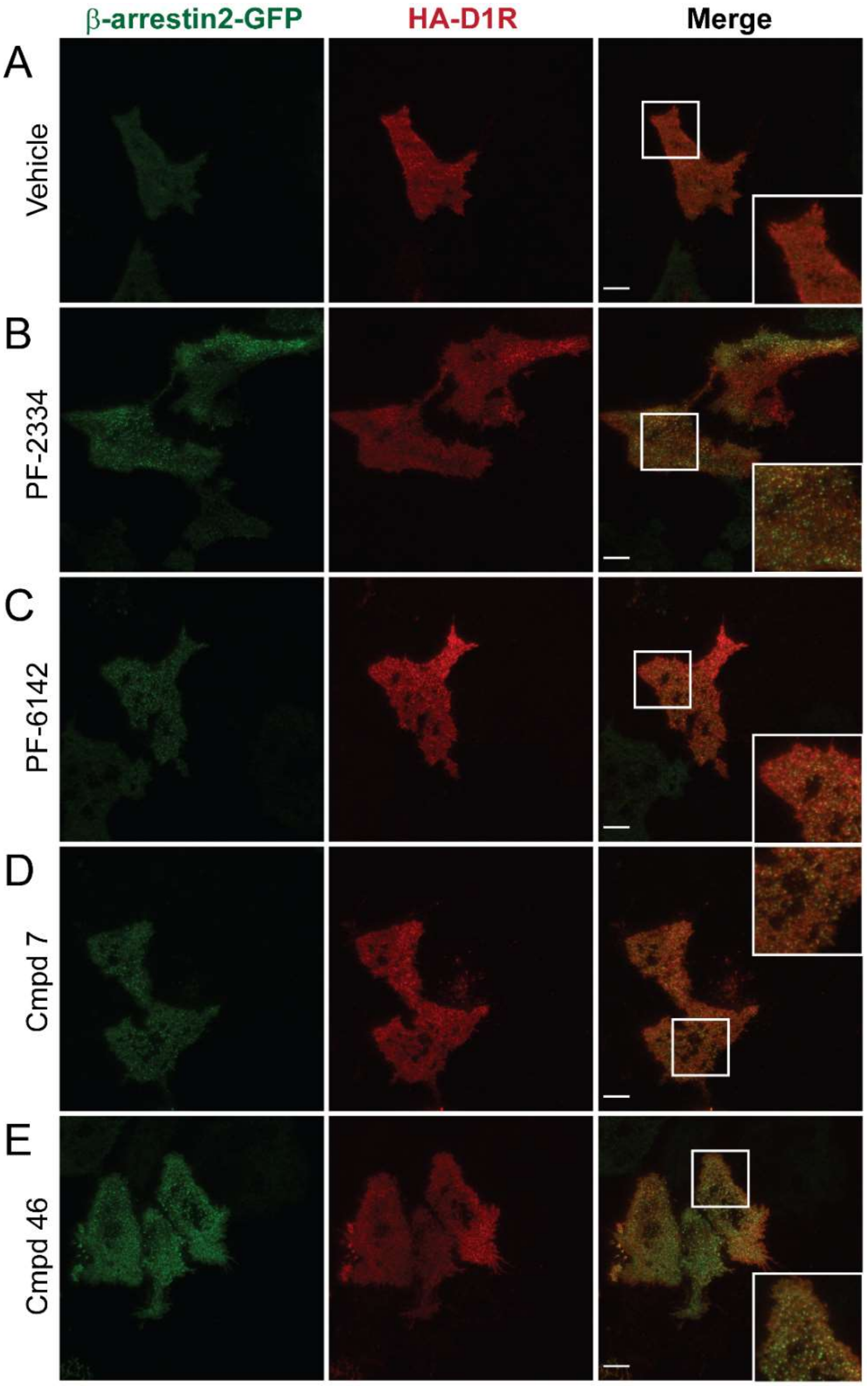


**Supplemental Figure 4. β-arrestin2-GFP recruitment to cell surface HA-D1R after treatment with non-catechol agonists.** HEK293 cells were transfected with HA-D1R and β-arrestin2-GFP. The cells were then treated with 3 uM of the indicated agonist and fixed after 10 minutes. The D1R was then detected using antibodies against the HA tag and visualized with alexa594 secondary antibodies. Punctate GFP signal indicates β-arrestin recruitment to the plasma membrane. TIRF microscopy was used to image a thin slice of the cells at the plasma membrane. **A)** Vehicle treated cells did not recruit β-arrestin to the plasma membrane. **B)** Treatment with PF-2334 induced moderate β-arrestin recruitment and colocalization with HA-

D1R. **C)** PF-6142 induced moderate β-arrestin recruitment to the plasma membrane. **D)** Cmpd 7 weakly recruited β-arrestin to the plasma membrane. **E)** Cmpd 46 moderately recruited β-arrestin to the plasma membrane. Similar results observed in >21 cells across three independent experiments. White boxes indicate the area that is enlarged in the corner of merged images. Scale bar = 10 μm.
